# SETD5 dysfunction in human astrocytes drives IL-6-mediated neuronal impairments via the JAK/STAT signaling pathway

**DOI:** 10.64898/2026.04.05.716613

**Authors:** Angels Almenar-Queralt, Isabella R. Fernandes, Janaina S. de Souza, Anju Singh, Xiwei Shan, Georgia Chaldaiopoulou, Dylan Phan, Kelsey Dang, Max Chang, Adriano Ferrasa, Noel Southall, Luisa B.V. Coelho, Livia Luz S. Nascimento, Brett Taylor, Miguel F. Tenreiro, Luisa F. Paredes-Brito, Raul Calvo, Eugin Destici, Pinar Mesci, Roberto H. Herai, Marc Ferrer, Allen Y. Wang, Ivan Garcia-Bassets, Christopher Benner, Juan J. Marugan, Alon Goren, Alysson R. Muotri

## Abstract

Intellectual disability (ID) and autism spectrum disorder (ASD) are neurodevelopmental conditions marked by lifelong impairments in cognitive, motor, and social functions. Hundreds of genetic variants have been linked to these disorders, including mutations in chromatin regulators such as the SET-domain-containing protein 5 (*SETD5*) gene. Most studies linking *SETD5* loss-of-function to ASD/ID have focused primarily on neurons. However, while SETD5 is highly expressed in astrocytes, its role in glia cells remains poorly understood. Here, we examine how dysfunction of *SETD5* in human-induced pluripotent stem cell (hiPSC)-derived astrocytes affects neuronal physiology. We show that SETD5-deficient astrocytes have increased levels of extracellular reactive oxygen species (ROS), glutamate, and interleukins-6 and 8 (IL-6 and IL-8). Elevated astrocytic IL-6 exerts a non-cell autonomous harmful effect on healthy neurons. Using SETD5-deficient astrocytes as a screening platform, we identify the JAK/STAT pathway as an upstream regulator of abnormal IL-6 accumulation associated with SETD5 dysfunction. Accordingly, pharmacological inhibition of JAK-STAT signaling restores extracellular IL-6 to basal levels and partially rescues astrocyte morphology and neuronal deficits. Collectively, these findings highlight the JAK/STAT pathway as a key regulator of SETD5-mediated astrocytic function and suggest its potential as a therapeutic target for astrocytic-driven neuronal impairments in ASD and ID.

## Introduction

Autism spectrum disorder (ASD) is a neurodevelopmental condition characterized by complex and heterogeneous etiology and symptomatology, affecting learning, communication, and cognitive and behavioral functioning. A large population-based study estimated the heritability of ASD to be approximately 80% (D et al. 2019). Extensive whole-exome and whole-genome sequencing efforts have identified hundreds of genetic variants strongly associated with ASD, highlighting convergent molecular and cellular pathways underlying its pathophysiology (Satterstrom et al. 2020; X. Zhou et al. 2022; Gogate et al. 2024; Kimura et al. 2022; Fu et al. 2022). These pathways primarily involve genes regulating synaptic function and neuronal communication, as well as genes involved in chromatin remodeling and transcription regulation.

While most ASD research has historically focused on neurons, growing evidence suggests that astrocytes also play a crucial role in ASD pathophysiology (M. Allen et al. 2022a; Xiong et al. 2023a; Vakilzadeh and Martinez-Cerdeño 2023; Gzielo and Nikiforuk 2021; Scuderi and Verkhratsky 2020; Martins et al. 2026). Astrocytes, the most abundant glial cell type in the brain, are essential for normal brain development. They contribute to neuronal survival, synapse formation and maturation, maintenance of brain homeostasis, and regulation of neural circuits (Farizatto and Baldwin 2023; N. J. Allen and Eroglu 2017). In addition, astrocytes regulate innate and adaptive immune responses in the central nervous system and are key modulators of neuroinflammatory pathways through the secretion of cytokines and chemokines (Fisher and Liddelow 2024). Astrocytes respond to injury or disease by adopting distinct reactive states, characterized by differences in gene expression, morphology, and functional properties (Escartin et al. 2021; Sofroniew 2020; Burda et al. 2022).

Inflammatory astrocyte reactivity has been implicated in the pathophysiology of brain injuries and multiple neurodegenerative and neuroinflammatory diseases (Heneka et al. 2015; Q. Wang et al. 2007; Hausmann 2003; Ponath et al. 2018). Notably, activated astrocytes and elevated levels of proinflammatory cytokines have been detected in brain tissues from individuals with ASD, suggesting their involvement in ASD-associated neurobiological alterations (Vakilzadeh and Martinez-Cerdeño 2023; Vargas et al. 2005). Supporting a direct astrocyte-neuron interplay in ASD, co-culturing mouse astrocytes from Fragile X or Rett syndrome mouse models with control neurons alters neuronal function (Jacobs et al. 2010; Ballas et al. 2009). Similarly, co-culturing of ASD human induced pluripotent stem cell (hiPSC)-derived astrocytes with control hiPSC-derived neurons leads to reduced neurite complexity and synaptic density (Russo et al. 2018). Using this system, we also identified interleukin-6 (IL-6) as a major astrocyte-derived mediator of neuronal dysfunction, which can be reversed by blocking IL-6 signaling (Russo et al. 2018). Notably, transplantation of ASD-hiPSC-derived astrocytes into healthy mouse brains was sufficient to induce repetitive behaviors, memory impairments, and reduced long-term potentiation (M. Allen et al. 2022a). These findings identify astrocytes as active contributors to ASD pathology and underscore their potential as experimental platforms for discovering novel pharmacological targets to alleviate behavioral and cognitive symptoms.

Among ASD-linked high-risk genes, *SETD5* (SET domain-containing protein 5; OMIM 615743) has been identified as a chromatin and transcription regulator associated with ASD and intellectual disability (ID) (Fernandes et al. 2018). Although SETD5 is expressed across all human brain cell types, most studies to date have focused on its role in neurons, primarily using *Setd5* haploinsufficient mouse models (Moore et al. 2019; Sessa et al. 2019; Deliu et al. 2018; Nakagawa et al. 2020). These studies indicate that *Setd5* haploinsufficiency disrupts neuronal network connectivity, reduces synaptic density and synaptic proteins, and impairs neurite outgrowth in cultured cortical neurons (Moore et al. 2019; Sessa et al. 2019). In addition, *Setd5* deficiency alters neuronal proliferation, affects the generation of early-born neurons, and leads to autistic-like behaviors in mice (Moore et al. 2019; Sessa et al. 2019; Deliu et al. 2018; Nakagawa et al. 2020). At the molecular level, SETD5 regulates transcription via interactions with polymerase-associated factor 1 (PAF1C), histone deacetylase 3 (HDAC3), and nuclear receptor co-repressor (NCoRs) components (Osipovich et al. 2016; Deliu et al. 2018; Nakagawa et al. 2020; Matsumura et al. 2021; Yu et al. 2017), as well as through direct or indirect modulation of histone modifications including H3K36me, H3K9me, H3K27ac, and H4K16ac (Sessa et al. 2019; Z. Wang et al. 2020; Matsumura et al. 2021; Nakagawa et al. 2020; Li et al. 2023). However, the impact of SETD5 loss-of-function on glial cells populations remains largely understudied.

Our previous work provided evidence supporting a role of SETD5 in ASD astrocytes: conditioned media from hiPSC-derived astrocytes of an ASD individual carrying a SETD5 SET-domain mutation exhibited elevated levels of extracellular IL-6 and, when co-cultured with control neurons, induced IL-6-dependent impairments in neuronal morphology and synapse number (Russo et al. 2018). Here, to confirm and expand the impact of *SETD5* dysfunction on astrocyte physiology and in turn on neuronal function, we used isogenic pairs of astrocytes derived from (i) hiPSC established from the severely affected individual with ASD/ID carrying the SETD5-SET domain mutation and their CRISPR corrected counterparts, and (ii) CRISPR/Cas9-engineered *SETD5* haploinsufficient hiPSC lines generated from a neurotypical individual. Using these systems, we demonstrate that *SETD5* dysfunction drives the astrocytic population towards a more reactive state, characterized by morphological and functional changes, including aberrant accumulation of IL-6, which, in turn, alters neuronal physiology in co-culture. Through targeted drug screening, we identified the Janus kinase/signal transducer and activator of transcription (JAK/STAT) pathway as an upstream regulator of IL-6 dysregulation induced by SETD5 dysfunction. Pharmacological inhibition of this pathway attenuated SETD5-dependent astrocytic abnormalities and partially restored neuronal deficits. Together, these findings confirm a role of SETD5 in astrocyte physiology and suggest that modulation of the JAK/STAT pathway as a new therapeutic strategy not only for individuals with SETD5-related ASD/ID but also for broader ASD populations exhibiting IL-6-driven neuroinflammation.

## Results

### SETD5 loss-of-function alters astrocytes’ morphology

In the human brain, *SETD5* is expressed in astrocytes at levels comparable to neurons, and higher to some neuronal subtypes (**Fig. 1a**). To evaluate the impact of SETD5 dysfunction on astrocytes, which to our knowledge, has not been thoroughly studies previously, we generated astrocytes from hiPSC-derived neuronal progenitor cells (NPCs) (**Supp. Fig. 1 and 2**) using a previously described differentiation protocol with minor modifications (**Supp. Fig. 3**) (Russo et al. 2018). Astrocytes derived from neurotypical hiPSCs exhibit higher *SETD5* levels than fibroblasts, hiPSCs, and hiPSC-derived NPCs, as well as hiPSC-derived neurons (**Fig. 1b** and **Suppl. Table 1 and Suppl. Table 2**).

**Figure 1.**
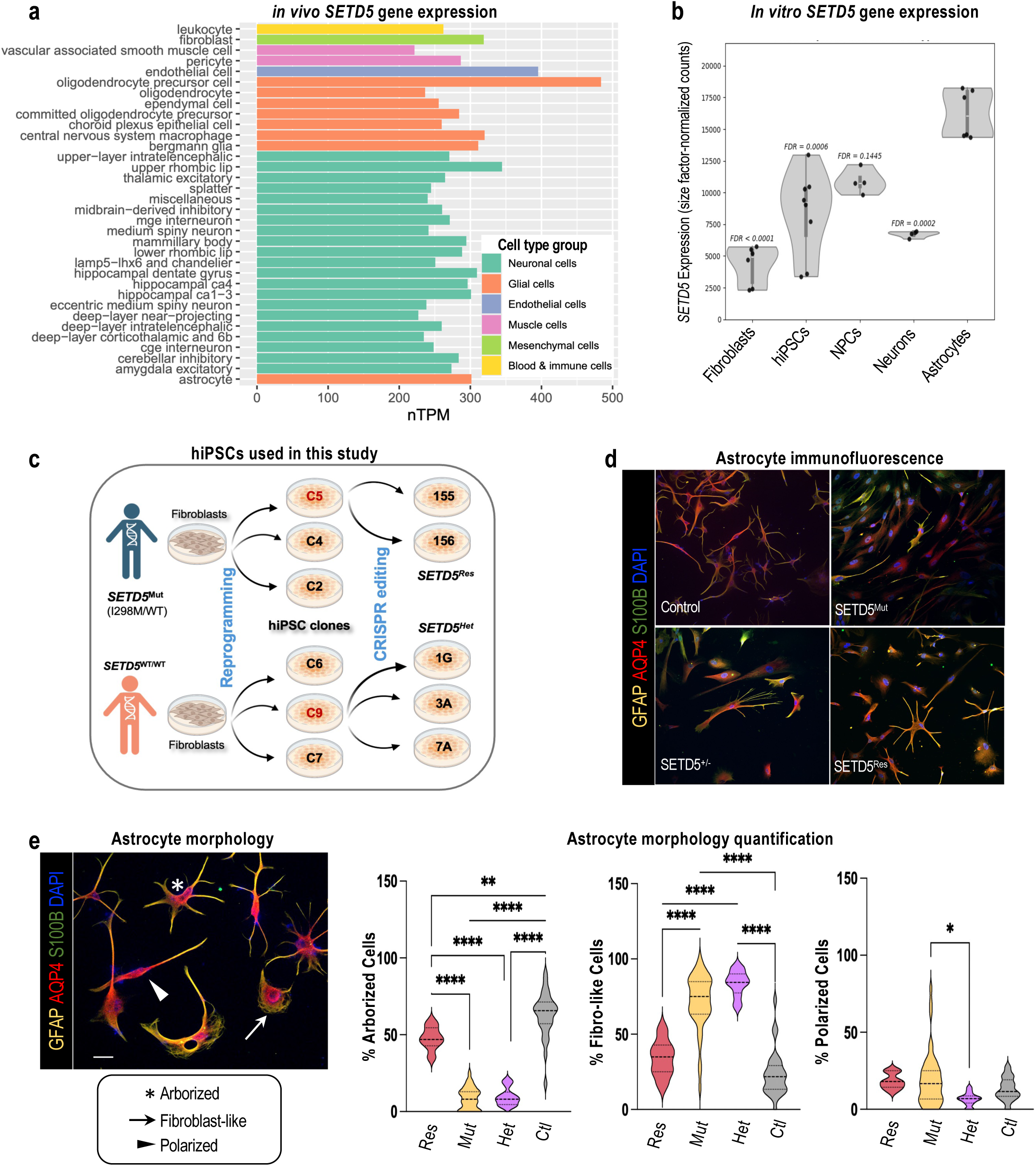
SETD5 dysfunction alters hiPSC-derived astrocytes morphology. **a.** Bar plot showing normalized transcript expression levels (nTPM) of *SETD5* across a range of annotated human brain cell types. Each bar represents a distinct cell type, grouped by major cell class as indicated by color coding. **b**. Violin plot representing the expression profile of the *SETD5* gene across human fibroblasts (n=6), hiPSCs (n=8), NPCs (n=4), neurons (n=4), and astrocytes (n=6). Biological replicates of each cell type are represented as black circles. Expression profiles of each biological replicate are represented as size-factor normalized counts (Suppl. Table 1). FDR (false discovery rate) statistical significance is relative to astrocytes (Suppl. Table 2). **c.** Schematic representation showing the generation and the different hiPSC clonal lines used in this study. Two different hiPSCs clones – one from an individual carrying a *SETD5* mutation (SETD5^I298M^) and one from a neurotypical individual – were previously established. In this study we report the subsequent CRISPR editing to correct the mutation from one of the SETD5^I298M^ clone, and the generation of heterozygous KO SETD5 lines from one of the control clones. *SETD5^I298M/WT^*(*SETD5^Mut^*), *SETD5^WT/WT^* (Control or *SETD5^Res^*), *SETD5^WT/KO^* (*SETD5^Het^*) **d.** Representative immunofluorescence images of control (*SETD5^WT/WT^)*, *SETD5^Mut^*, *SETD5^Het^*, and *SETD5^WT*/WT^* astrocytes stained for GFAP (yellow), AQP4 (red), S100ß (green), DAPI (blue), indicating morphological differences between the conditions. Scale bar 50 µm. **e.** Higher magnification representative image from control cells on the left illustrates the morphological distinctions, categorized as arborized (asterisk), polarized (arrowhead), and fibroblast-like (arrow). GFAP (yellow), AQP4 (red), S100ß (green), DAPI (blue). Scale bar 50 µm. Quantitative analysis of cell populations based on morphology across different genotypes: control (3 clones, 26 pictures, 396 astrocytes), *SETD5^Mut^* (2 clones, 33 pictures, 377 astrocytes), *SETD5^Het^* (2 clones, 14 pictures, 310 astrocytes), and *SETD5^Res^* (1 clone, 6 pictures, 83 astrocytes), showing the proportion of cells with each morphological phenotype. Ordinary one-way ANOVA, multiple comparisons Dunnett’s test.

**Figure 2.**
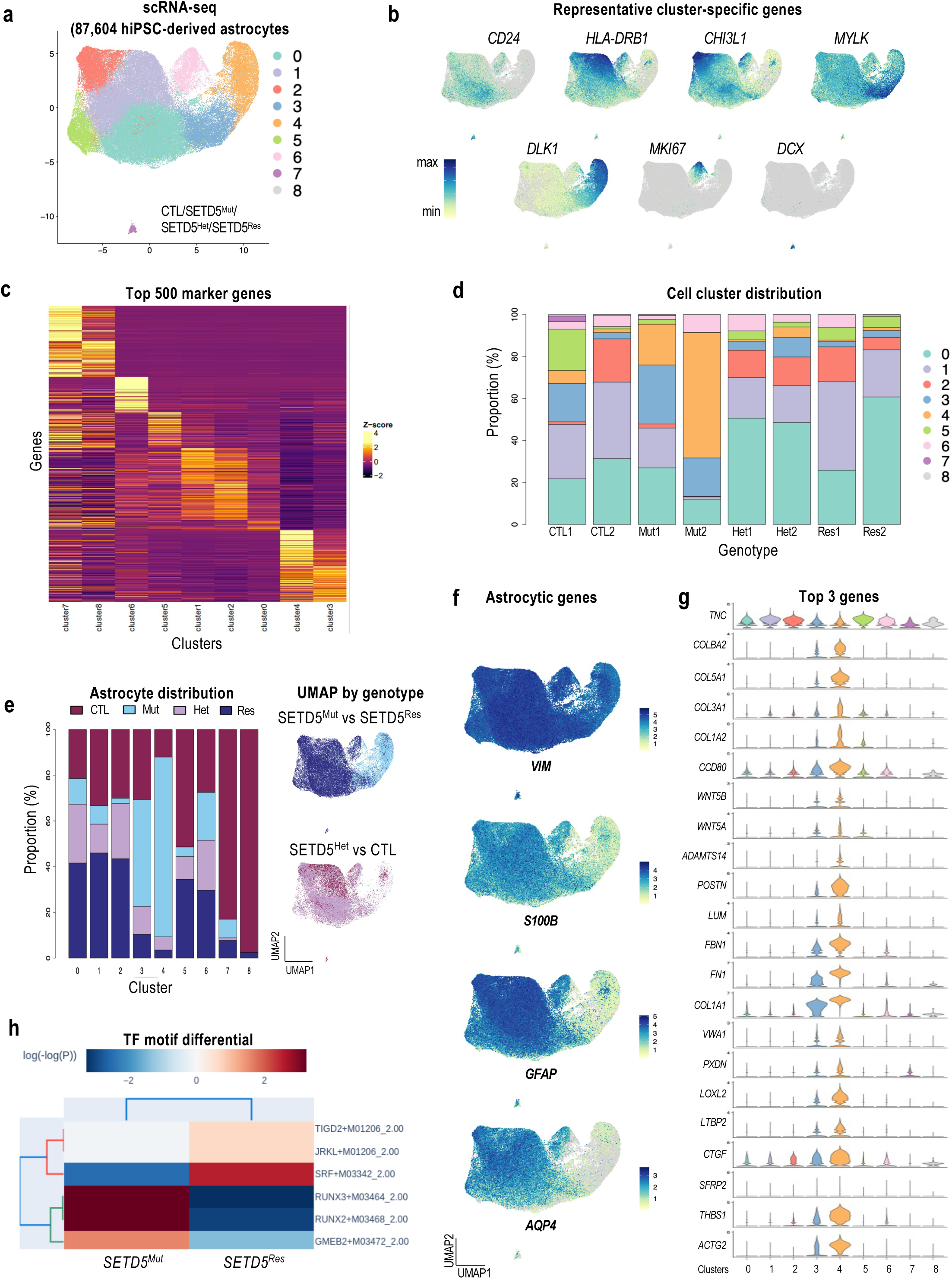
**single-cell transcriptome analysis enables identification of hiPS-derived astrocyte cell composition across the different lines**. **a.** Uniform Manifold Approximation and Projection (UMAP) plot displaying the distribution of scRNA-seq from 87,604 hiPSC-derived astrocytes, including Control, *SETD5^Mut^*, SETD5^Het^, and *SETD5^Res^*. The analysis identified distinct clusters with defined gene expression profiles. **b.** UMAP plots showing the spatial distribution of indicated genes across the dataset, with each point representing a single cell. Expression levels are color-coded from low (light yellow) to high (dark blue), as indicated by the colorbar. Grey=not expressed **c**. Heatmap displaying the top 500 marker genes across clusters (columns = clusters; rows = genes). Expression values are shown as Z-scored CPM. Marker genes were defined by differential expression (FDR < 0.05, avg log2FC > 0.58) and ranked by decreasing avg log 2FC. **d**. Bar graph showing the relative distribution of cell clusters within each genotype: reference (Control), mutant (*SETD5^Mut^*), heterozygous (*SETD5^Het^*), and rescue (*SETD5^Res^*) lines. **e.** Bar graph showing the relative distribution of astrocytes from each genotype, including Control, *SETD5^Mut^*, *SETD5^Het^*, and *SETD5^Res^* per cluster. On the right, a UMAP visualization of gene expression showing the distribution of cells colored by the indicated genotypes. **f.** UMAP plots showing the spatial distribution of indicated genes across the dataset, with each point representing a single cell. Expression levels are color-coded from low (light yellow) to high (dark blue), as indicated by the color bar. **g.** Violin plot of the top three genes expressed in each cluster. **h.** Heatmap of top motif enrichment from snATAC-seq of astrocytes derived from the isogenic hiPSCs pair *SETD5^Mut^* vs *SETD5^Res^*.

**Figure 3.**
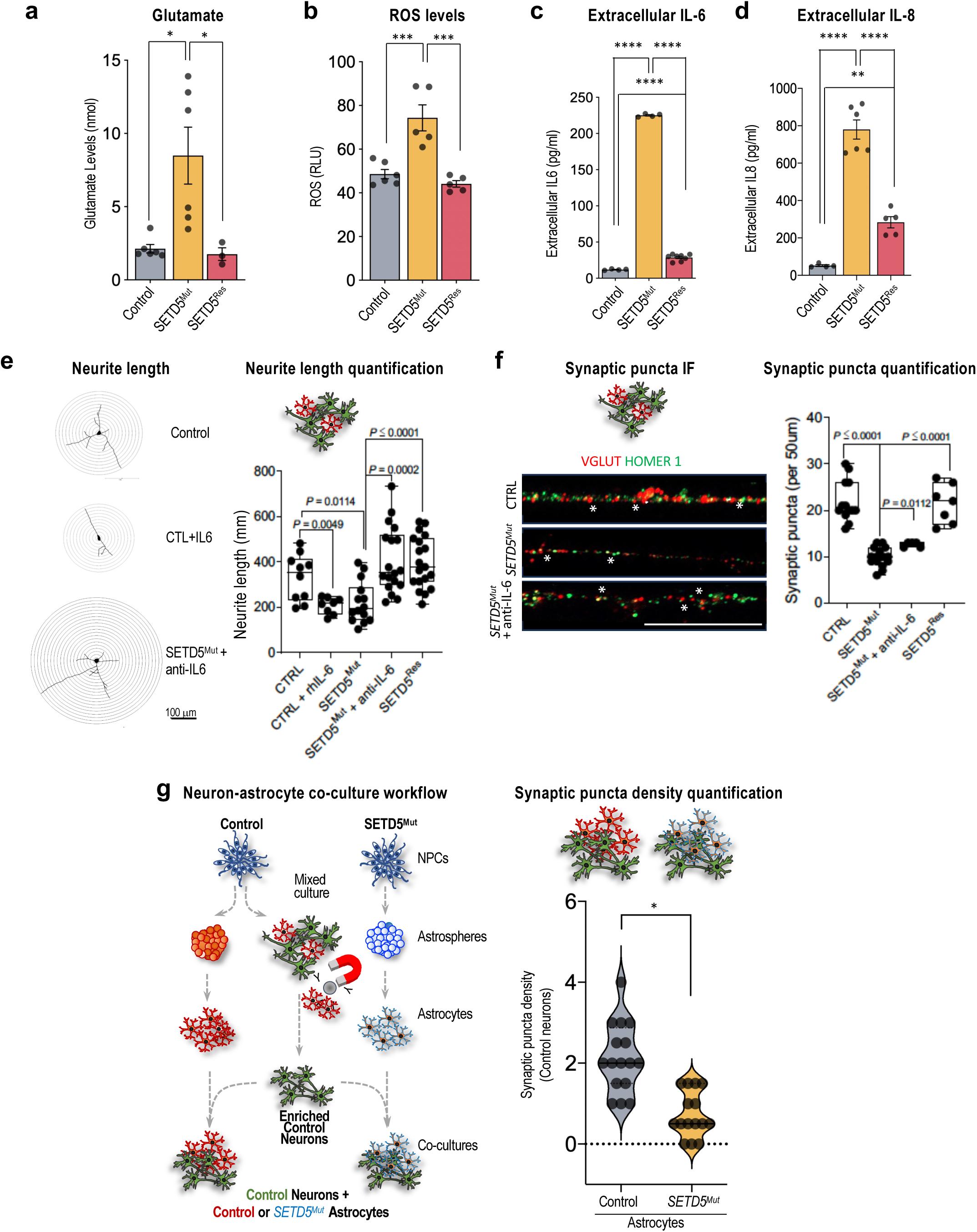
Functional and Morphological Assessment of Neurons with *SETD5* Mutation and IL-6 Inhibition. **a.** Bar graph showing glutamate uptake levels in control, *SETD5^Mut^*, and *SETD5^Res^* astrocytes. **b.** Reactive oxygen species (ROS) production is quantified in the same cell types. **c.** and **d.** results display elevated IL-6 and IL-8 levels secreted by *SETD5^Mut^* astrocytes. **e.** Graph comparing neurite length of hiPSC-derived neurons using Sholl analysis across indicated genotypes and treatments, accompanied by representative neurons with concentric circles (Sholl radii) centered on the soma, illustrating the method used to quantify neurite length. Measurements were done in non-enriched neuronal cultures. Each dot represents an individual cell, with bars indicating mean ± SEM. Right. Scale bar: 100 μm. **f.** Representative immunofluorescence images showing staining of non-enriched hiPSC-derived neuronal cultures from indicated genotypes with the pre-synaptic marker VGLUT and the post-synaptic marker HOMER1 on MAP2+ (not shown) neurites. Asterisks mark examples of synaptic puncta (Right). Graphic representation of the synaptic puncta in control, *SETD5^Mut^*, *SETD5^Mut^* + anti-IL-6, and *SETD5^Res^* neurons. Ordinary one-way ANOVA, multiple comparisons Dunnett’s test. P value < 0.05. **g.** Left: Illustration of the procedure used for the generation of neuronal-enriched cultures and the neuron-astrocyte co-culture system. Right: violin plots of synaptic puncta density in control-enriched neurons co-cultured with either control or *SETD5^Mut^* astrocytes.

Following this protocol, we derived astrocytes from: (i) hiPSCs generated from a severely intellectually affected individual carrying a point mutation in the SET-domain of *SETD5* (NM_001080517: exon9: c.A894G: p.I298M, hereafter referred to as *SETD5^Mut^*) (Russo et al. 2018); (ii) CRISPR-corrected *SETD5^Mut^* hiPSCs to confirm mutation specificity hereafter referred to as SETD5 Rescue (*SETD5^Res^*); (iii) previously characterized hiPSCs derived from a neurotypical individual (hereafter referred to as control or reference) (Marchetto et al. 2017); and (iv) CRISPR-engineered *SETD5* haploinsufficient hiPSCs, hereafter referred as *SETD5^Het^* (**Fig. 1c** and **Supp. Fig. 1**).

As expected, all these hiPSC-derived astrocytic populations stained for canonical astrocytic markers, including glial fibrillary acidic protein (GFAP), S100 Calcium-binding protein B (S100B), and Aquaporin 4 (AQP4) (**Fig. 1d**, **Supp. Fig. 3**). Remarkably, however, we observed evident morphological differences in *SETD5^Mut^* and *SETD5^Het^* astrocytes when compared to *SETD5^Res^* and controls (**Fig. 1d-e**). Following the same morphological categories as previously described (Jones et al. 2017), control astrocytes were predominantly arborized, with long, thin, branched processes (>2 per cell), alongside smaller populations of polarized (thin, elongated) and fibroblast-like (process-devoid) cells (**Fig. 1d-e**). In contrast, *SETD5^Mut^*astrocyte cultures were less diverse and mostly fibroblast-like, with a marked reduction in arborized cells (**Fig. 1d-e**). This phenotype was largely reverted in *SETD5^Res^*astrocytes, which resembled controls (**Fig. 1d-e**). *SETD5^Het^* cells also consisted mainly of fibroblast-like astrocytes, with a statistically loss of arborized and polarized populations, closely resembling *SETD5^Mut^*(**Fig. 1d-e**); aside from a slightly higher proportion of polarized cells in *SETD5^Mut^*, the two genotypes were otherwise similar (**Fig. 1d-e**). Together, these results indicate that SETD5 dysfunction—whether through a missense mutation or heterozygous loss—does not prevent astrocyte differentiation from NPCs but shifts the populations toward fibroblast-like morphologies.

### SETD5 loss-of-function alters astrocyte cellular populations

To investigate whether the SETD5-dependent morphological differences reflect distinct astrocyte types or states, we performed single-cell RNA-seq (scRNA-seq) on astrocytes derived from two hiPSC clones per genotype to a total of 87,604 cells profiles. Joint analysis of the merged scRNA-seq dataset identified nine cell clusters (**Fig. 2a**); each defined by a unique transcriptional profile, with representative marker gene expression shown across the UMAP projection (**Fig. 2a, b**; top 2-3 cluster-enriched genes are displayed in **Supp. Fig. 4a**). Clusters 0 and 1 showed notable expression of immune-related genes, such as *CD24* and various HLA genes (**Fig. 2b**). Cluster 2 was defined by genes like *CHI3L1*, a marker for neuroinflammation (Mwale et al. 2025) (**Fig. 2b**). Clusters 3 and 4 show enrichment for *FN1* and *MYLK*, respectively, (**Fig. 2b**) consistent with a mesenchymal-like, contractile reactive astrocyte state associated with injury and tissue remodeling (O’Shea et al. 2024; Anderson et al. 2016; Wiese et al. 2012; Sofroniew 2009, 2020; Hemati-Gourabi et al. 2025; Hara et al. 2017; Shih et al. 2014). Cluster 5 was represented by *DLK1*, previously linked to injury-responsive glial cells (Jin and Zheng 2019); cluster 6 was enriched in markers such as *TOP2A* and *MKI67*, commonly used to identify dividing cells (Spathopoulou et al. 2024); clusters 7 and 8 included neuronal genes, such as *DCX*, a canonical marker of immature neurons and neuronal progenitors, consistent with these clusters representing early neuronal lineage cells (Karl et al. 2005) **(Fig. 2a-b**; **Supp. Fig. 4a**, and **Suppl. Table 3**). Supporting the robustness of our astrocyte differentiation protocol, these neuronally enriched cluster 7/8 cells represented no more than 0.59% of total cells in any genotype and were mainly contributed by one of the control samples (**Fig. 2c,d**).

**Figure 4.**
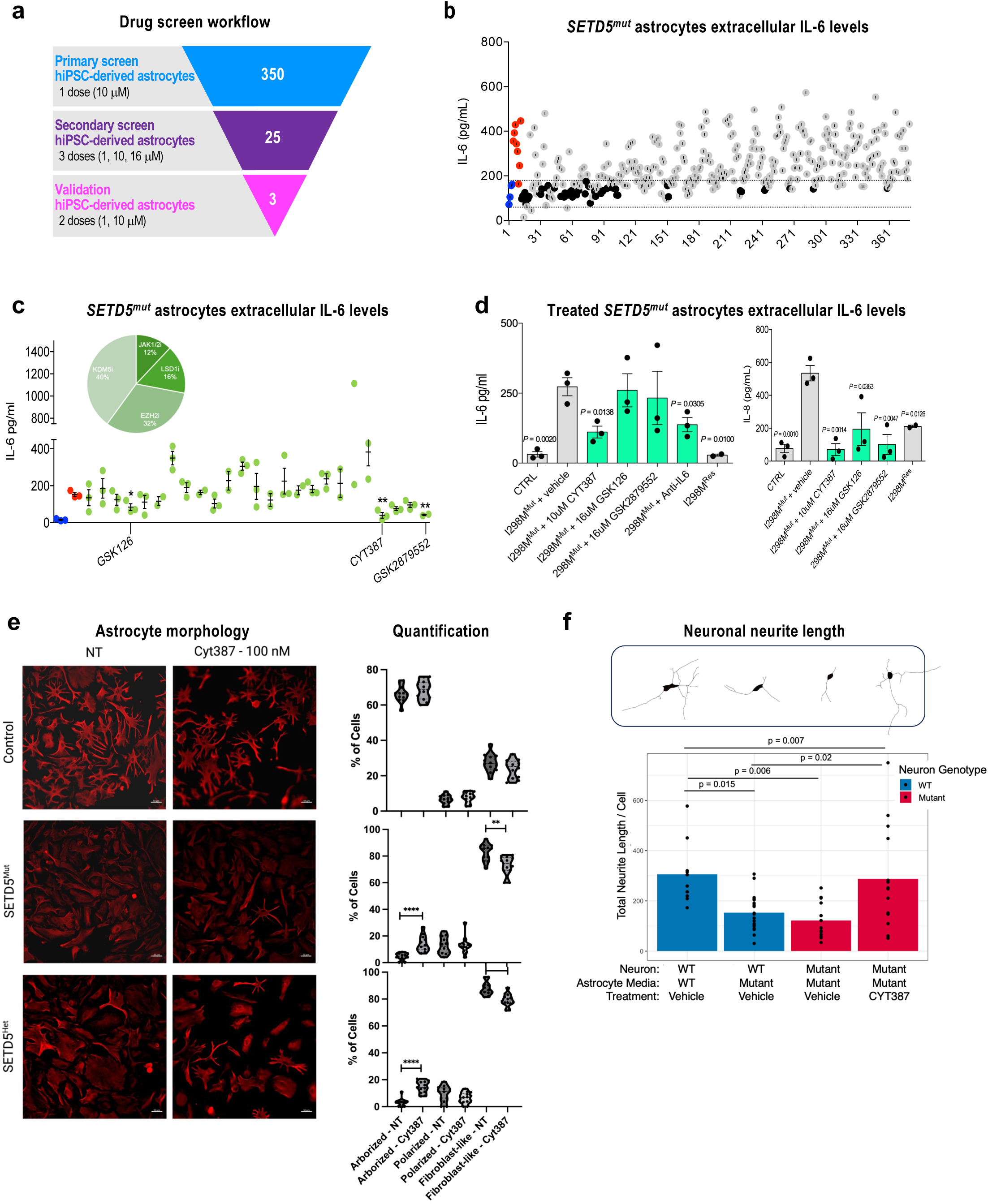
Pharmacological Modulation of IL-6 Levels in *SETD5^Mut^* Astrocytes and Neural Response. **a.** Schematic depicting the screening process using *SETD5^Mut^*astrocytes to identify compounds that reduce IL-6 levels and maintain cell viability, leading to the discovery of the JAK/STAT pathway as an upstream regulator of the accumulation of extracellular IL-6 in *SETD5^Mut^* astrocytes. **b.** High-throughput screening results showing the IL-6 levels after treatment with a 350-compound library, leading to 25 hits, reducing IL-6 levels without affecting cell viability. **c.** Graph representing levels of extracellular IL-6 after treating *SETD5^Mut^* astrocytes with 25 selected compounds, accompanied by a pie chart showing the distribution of inhibitors categories comprised in the selected hits. **d.** Graph representing levels of extracellular IL-6 (left) and IL-8 (right) in hiPSC-derived astrocytes of the indicated conditions. **e.** Left: representative immunocytochemistry images. Right: quantitative analysis of hiPSC-derived astrocyte morphology categories from indicated genotypes treated with vehicle or CY387. **f.** Graph representing neurite length in the indicated conditions. Representative morphology of reconstructed hiPSC-derived neurons expressing GFP-synapsin using Neurolocida software placed on top of the graph in the corresponding condition.

Importantly, comparing the proportion of each genotype within clusters revealed that *SETD5^Mut^* astrocytes were the primary contributors to clusters 3 and 4 relative to other samples (**Fig. 2c, d**). While *SETD5^Het^* astrocytes also contributed to these clusters, they were represented similarly to their isogenic controls (**Fig. 2c**) and contributed to other clusters in comparable proportions (**Fig. 2d**). Closer examination of these two clusters reveals that while these cells express glial markers Vimentin (*VIM*) and *S100B* (like the rest of the clusters), they contain substantially reduced number of cells expressing the classical astrocytic markers *GFAP* and *AQP4* (**Fig. 2e**). Clusters 3 and 4 astrocytes express genes encoding secreted extracellular matrix (ECM) components, including *COL8A2*, *COL5A1*, *COL3A1*, *COL1A1*, *CAL1A2*, *FBN1*, and *PSTN* (**Fig. 2f**). In contrast, cells expressing *TNC* (Tenascin C), another ECM glycoprotein are found at comparable levels across all clusters (**Fig. 2f**).

Gene Ontology (GO) analysis highlighted ECM organization among the top categories in cluster 4, the most enriched in *SETD5^Mut^* astrocytes suggesting that these mutant astrocytes may adopt a reactive transcriptional program (**Fig. 2f**) (O’Shea et al. 2024; Anderson et al. 2016; Wiese et al. 2012; Sofroniew 2009, 2020; Hemati-Gourabi et al. 2025; Hara et al. 2017; Shih et al. 2014).

We further supported the distinct transcriptional status of *SETD5^Mut^*astrocytes by single-nucleus (sn)ATAC-seq on these cells and its isogenic *SETD5^Res^*. A heatmap of the most significantly enriched DNA binding motifs for transcription factors in *SETD5^Mut^* astrocytes revealed enrichment in the RUNX2/3 and GMEB2 motifs while depletion in the SRF motif (**Fig. 2g**). RUNX2/3 motif enrichment in *SETD5^Mut^* astrocytes may reflect alterations in the differentiation state or activation of reactive/inflammatory programs in mutant astrocytes (Lu et al. 2024; Shen et al. 2025). RUNX2 is a well-known regulator of ECM genes in multiple cell types (Shen et al. 2025), suggesting that it may directly regulate ECM genes in *SETD5^Mut^* astrocytes. Although the role of the transcription factor GEMB2 in astrocytes has not been studied, GMEB2 motif enrichment could be linked to stress or injury-responsive gene programs regulation, as GMEB2 belongs to a family of transcription modulators that interact with the glucocorticoid receptor and influence glucocorticoid-regulated transcriptional responses (Kaul et al. 2000). Reduced accessibility at SRF motifs may indicate alterations in transcriptional regulation linked to astrocyte dynamic morphological changes and reactivity, consistent with studies showing that SRF is critical for maintaining astrocytes in a non-reactive state and that its loss leads to reactive-like phenotypes (Jain et al. 2021). Motif enrichment analysis revealed that RUNX2/3- and SRF-associated differentially accessible peaks map to a large number of unique protein-coding genes, consistent with broad transcriptional regulatory effects (**Supp. Fig. 5a)**. Pathway enrichment of these motif-associated regions highlighted processes related to ECM organization, cell adhesion, and wound response, reinforcing a shift toward a reactive and remodeling phenotype (**Supp. Fig. 5b)**. Integration with gene expression data showed that several putative SRF and RUNX3 target genes exhibit concordant changes in chromatin accessibility and transcription, supporting functional coupling between motif accessibility and downstream gene regulation (**Supp. Fig. 5c)**. Finally, inferred TF-gene regulatory interactions revealed that RUNX2 and RUNX3 were connected to a substantially larger number of predicted target genes than SRF and GMEB2, (**Supp. Fig. 5d)**. Given the high similarity between RUNX2 and RUNX3 binding motifs, this shared connectivity likely reflects, at least in part, an inability to distinguish the two factors’ binding sites rather than confirmed independent or coordinated co-regulation. Nonetheless, the individual breadth of RUNX2- and RUNX3-associated peaks (Suppl. Fig. 5a) points to RUNX family motifs as broadly represented across the altered chromatin landscape in *SETD5^Mut^* astrocytes. Together, these data suggest that the *SETD5* mutation biases astrocytes toward a stress-responsive, ECM-remodeling chromatin state.

### IL-6-mediated neurotoxicity originates from SETD5-mutant astrocytes

Reactive astrocytes are characterized by excessive glutamate release, oxidative stress, and increased secretion of pro-inflammatory cytokines (Ben Haim et al. 2015). To determine whether the transcriptional and epigenomic changes observed in *SETD5^Mut^*astrocytic populations lead to transitioning into a reactive astrocyte state, we measured extracellular glutamate, reactive oxidative species (ROS), and the pro-inflammatory cytokines interleukin-6 (IL-6) and interleukin-8 (IL-8) in control, *SETD5^Mut^*, and *SETD5^Res^* hiPSC-derived astrocytes. We found that all these factors were elevated in the conditioned media of *SETD5^Mut^* astrocytes and were substantially reduced in *SETD5^Res^* astrocytes (**Fig. 3a-d**), indicating that a missense mutation in the region encoding for the SET domain of SETD5 is sufficient to drive astrocyte reactivity. Supporting the contribution of a loss-of-function mechanism, we also observed increased IL-6 levels in the conditioned media of *SETD5^Het^* astrocytes compared with isogenic controls, though to a lesser extent than the media from *SETD5^Mut^*astrocytes (**Suppl. Fig. 6**).

Elevated extracellular IL-6 in hiPSC-derived astrocyte-neuron co-cultures adversely affects neuronal morphology and function, including in models of Parkinson’s disease and ASD (Pons-Espinal et al. 2024; Russo et al. 2018; Goshi et al. 2025). To investigate the impact of the IL-6 pathway in *SETD5^Mut^*neural cultures, we differentiated control and *SETD5^Mut^* NPCs into neurons, which may contain a low percentage of astrocyte-like cells as previously described (Russo et al. 2018). *SETD5^Mut^* neurons show decreased neurite length in a manner that is similar to control neurons treated with recombinant human IL-6 (rhIL-6) (**Fig. 3e**). Importantly, the *SETD5^Mut^* neuronal phenotype can be rescued by reversing the missense mutation in the SET domain in *SETD5^Res^* neurons (**Fig. 3e**). Furthermore, *SETD5^Mut^* neurons treated with IL-6 blocking antibodies (anti-IL-6) restored dendrite length to control and *SETD5^Res^* levels (**Fig. 3e**), demonstrating that neutralizing IL-6 activity can mitigate its detrimental effects on neuronal morphology, consistent with previous findings (Russo et al. 2018). Similarly, the reduced number of synaptic puncta per μm observed in *SETD5^Mut^* neurons compared with controls and *SETD5^Res^*neurons was partially restored after incubation with anti-IL-6 (**Fig. 3f**). Together, these results confirm the involvement of SETD5 in maintaining astrocytic and neuronal physiology, and suggest that *SETD5^Mut^* neuronal defects are caused by IL-6 that we postulated could be generated by residual *SETD5^Mut^* astrocytes or precursors generated during the process of neuronal differentiation.

To confirm the contribution of *SETD5^Mut^* astrocytes to neuronal morphological alterations, we established neuron-astrocyte co-cultures as previously reported (Russo et al., 2018) (**Fig. 3g**). To first deplete residual astrocytes that might be generated during derivation of neuronal cultures, we used magnetic beads coated with antibodies targeting the astrocytic marker CD44 to eliminate this fraction (**Fig. 3g**). The resulting control neurons were then co-cultured with either control or *SETD5^Mut^*astrocytes derived from corresponding NPCs (**Fig. 3g**). We observed a reduction in synaptic puncta density, quantified by co-staining of the presynaptic marker vesicular glutamate transporter 1 (VGLUT1) and the postsynaptic marker HOMER1, in control neurons co-cultured with *SETD5^Mut^* astrocytes compared with those co-cultured with control astrocytes (**Fig. 3g**). Together, these results confirm that the missense mutation in the SET domain of SETD5 is sufficient to lead to an astrocyte reactive state that negatively impacts neuronal morphology, with IL-6 playing a key role in these defects.

### Drug screening identifies JAK/STAT inhibition as a strategy to reduce IL-6 and neuronal defects driven by *SETD5****^Mut^*** astrocytes

Elevated IL-6 has been previously linked to autistic-like behavior in maternal immune activation mouse models (Smith et al. 2007), and has also been consistently observed in the brains of individuals with ASD (H. Wei et al. 2013). Overexpression of IL-6 in mice induces ASD-like behaviors and neuronal morphological changes (Hongen Wei et al. 2012), highlighting its role in neurodevelopmental pathology. Neuroinflammation is increasingly recognized as a key contributor to ASD and other neurodevelopmental disorders (Xiong et al. 2023b), and astrocytes are critical modulators of this process, influencing cytokine release, synaptic regulation, and neuroimmune interactions relevant to ASD pathology (M. Allen et al. 2022b). Based on these observations and our findings that an ASD/ID-linked *SETD5* mutation drives astrocytic IL-6-dependent neuronal defects, targeting astrocytic IL-6 represents a promising therapeutic strategy.

To explore a pharmacological approach, we used *SETD5^Mut^*-derived astrocytes as a platform to identify compounds capable of reducing extracellular IL-6 produced (**Fig. 4a**). Given the potential role of SETD5 as a lysine methyltransferase that influences transcription, chromatin structure, and RNA processing (Li et al. 2023), we screened an epigenetic library of 350 compounds (**Suppl. Table 4**). From this primary screening, 25 compounds were selected based on their ability to reduce extracellular IL-6 levels while showing low impact on cell viability (**Fig. 4c**). Although not used to select hit compounds, their effect on extracellular IL-8 levels was assessed in parallel to evaluate broader impact on cytokine signaling (**Suppl. Fig. 6**). Among the compounds reducing extracellular IL-6 levels, 40% are lysine-specific demethylase-5 (KDM5, also known as JARID1) inhibitors; 32% are histone methyltransferase Enhancer of Zeste Homolog 2 (EZH2) inhibitors; 16% are Lysine-Specific Demethylase-1 (LSD1) inhibitors; and 12% are Janus Kinase 1/2 (JAK1/2) inhibitors (**Fig. 4c**). After a second validation of these 25 compounds in independent *SETD5*^Mut^ astrocyte cultures at 2 different concentrations, we identified three compounds that lowered extracellular levels of IL-6 without impacting viability (**Fig. 4c**). Specifically, GSK126, a potent and highly selective small-molecule inhibitor of the enzyme EZH2 (Enhancer of Zeste homolog 2), a key component of the Polycomb Repressive Complex 2 (PRC2) complex; GSK2879552 is a potent and irreversible inhibitor of lysine-specific demethylase 1 (LSD1 or KDM1A); and CYT387 (also known as momelotinib), which targets and blocks the activity of Janus Kinase inhibitor 1 and 2 (JAK1/2) inhibiting the JAK-STAT signaling (**Fig. 4c**). These hits were assessed in independent *SETD5*^Mut^ astrocyte cultures, which led to narrow down to CYT387, which consistently reduced IL-6 levels (**Fig. 4e**).

To further evaluate the involvement of JAK1/2 or other JAK members, we assessed the ability to reduce extracellular IL-6 levels in independent batches of cultured *SETD5^Mut^* astrocytes with 24 new compounds comprising other JAK1, JAK2, or JAK3 inhibitors, and a few non-JAK inhibitors. While some compounds potently reduced IL-6 levels in a dose dependent way (**Suppl Fig. 6b**; green arrowheads), most, except CYT387 (Momelotinib) and BM-911543, reduced cell viability at higher concentrations (**Suppl Fig. 6b**). CYT387 also efficiently reduced extracellular IL-6 levels from *SETD5*^Het^ astrocytes (**Suppl Fig. 6c).**

Next, we examined whether CYT387 could restore the altered morphology of *SETD5^Mut^* astrocytes. Control, *SETD5^Mut^*, and *SETD5^Het^* were incubated with CYT387, and morphological analysis indicated that the compound partially restored astrocytic morphology by significantly decreasing the proportion of fibroblast-like cells while increasing the proportion of arborized cells (**Fig. 4e**). Notably, CYT387 did not change the relative proportions of the distinct morphological types in control cells (**Fig. 4e**). Finally, we tested whether CYT387 could prevent neuronal defects caused by S*ETD5^Mut^*-derived astrocytes. *SETD5^Mut^* neurons incubated with conditioned media from CYT387-treated S*ETD5^Mut^*-astrocytes showed restored total neurite length compared to *SETD5^Mut^* neurons exposed to media from vehicle-treated S*ETD5^Mut^*-astrocytes (**Fig. 4f**). Consistent with previous observations, control neurons exposed to conditioned media from vehicle-treated S*ETD5^Mut^*-astrocytes exhibited reduced neurite length compared to those incubated by media from vehicle-treated control astrocytes (**Fig. 4f**). Together, these findings highlight the therapeutic potential of blocking JAK1/2 by CYT387 to mitigate *SETD5^Mut^* -induced morphological disruptions in both astrocytes and neurons.

## Discussion

In this study, we demonstrate that SETD5 loss-of-function significantly alters human astrocyte physiology, thereby disrupting neuronal function. Using isogenic hiPSC-derived astrocytes carrying either an ASD/ID patient-derived *SETD5* mutation or engineered SETD5 haploinsufficiency, we found that SETD5 dysfunction shifts astrocytic populations toward a reactive-like state, characterized by morphological changes, distinct transcriptional and epigenomic signatures, and abnormal accumulation of pro-inflammatory mediators such as IL-6. Astrocyte-derived IL-6 promotes neuronal alterations, suggesting a non-cell autonomous mechanism that acts on neuronal morphology and synaptogenesis. A targeted drug screen identified the JAK1/2 inhibitor CYT387 as an effective compound for reducing IL-6 levels in *SETD5*-deficient astrocytes, thereby normalizing astrocytic morphology and restoring neuronal integrity to a certain extent. Our data highlight the JAK/STAT pathway as a potential therapeutic target for SETD5-related ASD/ID and for broader IL-6-mediated neuroinflammation.

We previously demonstrated that control neurons co-cultured with astrocytes from individuals with ASD, including the one studied here carrying the *SETD5^I298M^* mutation, exhibited altered morphology and reduced synaptic puncta, which were reversed by IL-6 blockade (Russo et al. 2018). Here, we expand these findings using CRISPR-corrected *SETD5^I298M^* hiPSCs, showing that a missense mutation in the SET domain of SETD5 is sufficient to alter astrocytic function and, non-cell autonomously, disrupt neurite complexity and synapse number via IL-6. Moreover, CRISPR-engineered *SETD5* haploinsufficient astrocytes exhibited similar defects, indicating that proper levels of functional SETD5 are required to maintain astrocytic homeostasis and balanced neuron-astrocyte interactions. Overall, *SETD5* dysfunction leads to distinct astrocyte populations characterized by a more fibroblast-like phenotype, transcriptional changes, and increased accumulation of IL-6, glutamate, ROS, and other cytokines, such as IL-8.

Single-cell RNA sequencing revealed that SETD5 dysfunction shifts the astrocyte population. However, whether this effect emerges only during astrocytic differentiation or it arises earlier at the precursors NPCs is not addressed here. While SETD5-mutant or haploinsufficient NPCs showed no gross morphological differences by Nestin staining, undetected altered NPC physiology due to SETD5 dysfunction could contribute to skewed astrocytic differentiation. Indeed, top GO pathways associated with genes defining clusters 3 and 4, enriched with *SETD5^Mu^*^t^-astrocytes, highlighted ECM organization pathways are also identified as one of the top GO pathways upregulated in *Setd5*-knockout mouse embryonic stem cells (mESCs) (Osipovich et al., 2016), suggesting a broader role for SETD5 in other cell types including pluripotent stem cells.

In exploring transcriptional drivers of *SETD5^Mu^*^t^-astrocytes identity, we found enrichment of *CREB3L1* (cAMP responsive element-binding protein 3-like 1) also called OASIS (Old Astrocyte Specifically Induced Substance), expressing astrocytes in clusters 3 and 4 (**Suppl. Fig. 3**). CREB3L1 was previously shown to be a putative endoplasmic reticulum (ER) stress sensor likely contributing to the resistance of astrocytes to ER stress (Saito et al. 2007). CREB3L1 is embedded in the ER membrane and, upon ER stress or other stimuli, is transported to the Golgi apparatus, cleaved, and translocated to the nuclei to activate transcription of its gene targets, including ECM-related genes (Vellanki et al. 2010). Clusters 3 and 4 were enriched for collagen-related genes consistent with CREB3L activity (**Fig. 2**). Although its role in brain remains understudied *Creb3l1*-KO mice show delayed astrocyte differentiation, and CREB3L1 is upregulated in reactive astrocytes after brain injury, particularly near the injury sites, contributing to glial scar formation and impaired axonal regeneration (Iseki et al. 2012). Seizures, a common feature in ASD with 20-40% of individuals affected (Amiet et al. 2008; Besag and Vasey 2021; Tuchman and Rapin 2002), can induce reactive astrocytes (Sumadewi et al. 2024; Robel et al. 2015), and neuroinflammatory responses including cytokine release such as IL-6 (Lehtimäki et al. 2003; Vezzani et al. 2008), as well as ECM remodeling (Blondiaux et al. 2023), consistent with a bidirectional feed-forward loop between seizure activity and glial dysfunction. Together, these observations suggest that in SETD5 syndrome, astrocytes may have heightened CREB3L1 activity, which could exacerbate ECM deposition and inflammatory responses after seizures, potentially impairing neuronal recovery.

Our snATAC-seq data show increased accessibility of RUNX2-binding motifs in SETD5^Mut^ astrocytes, pointing to activation of RUNX2-dependent transcriptional programs. While RUNX2 is best known for its role in osteogenesis, recent work demonstrates that its knockdown attenuates neuroinflammation and glial activation, supporting a broader function in regulating inflammatory gene expression (Shen et al. 2025). These observations raise the possibility that aberrant RUNX2 activity contributes to the pro-inflammatory astrocyte state observed upon SETD5 loss. More broadly, our findings suggest that SETD5 may restrain neuroinflammatory programs by limiting accessibility to transcription factors such as RUNX2, a mechanism that warrant further investigations.

Growing evidence further reinforces that astrocytes are not passive bystanders but active contributors to ASD (Vakilzadeh and Martinez-Cerdeño 2023). In particular, while circulating IL-6 has been shown to promote proliferation of adult forebrain neuronal stem cells in mice (Storer et al. 2018), it has been repeatedly reported to be elevated in the blood and brain of individuals with ASD, and often correlates with the severity of behavioral symptoms (Zhao et al. 2021). Further, experimental elevation of brain IL-6 in mice recapitulates autism-like features (Hongen Wei et al. 2012). Consequently, targeting reactive astrocytes as a platform to elucidate modulators of inflammatory states has become a significant area of interest (Leng et al. 2022; Clayton et al. 2024).

Given that *SETD5^Mut^* astrocytes accumulate extracellular IL-6, they offer a pathophysiologic link and a model system well-suited for screening compounds to reduce IL-6. Through a targeted screen, we identified the JAK1/2 inhibitor CYT387 as a potent suppressor of IL-6 aberrant accumulation, which also partially restored SETD5-deficiency-induced astrocytic morphological changes and non-cell-autonomous effects on neuronal morphology. Similar effects with other JAK inhibitors highlight the JAK/STAT3 pathway as a key upstream regulator of IL-6 aberrant accumulation in SETD5-deficient astrocytes (**Suppl. Fig. 6**) and point to this signaling axis as a promising therapeutic target for ASD and related conditions involving astrocyte-mediated IL-6 neuroinflammation. In support of these findings, a recent report using pooled CRISPRi screening in hiPSC-derived astrocytes also uncovered STAT3 as a key modulator of IL-6 astrocytic production, and further validated this observation *in vivo* using *Stat3*-KO mice (Leng et al. 2022). Multiple studies have implicated JAK/STAT signaling in ASD or ASD-model systems (Khera, Mehan, Kumar, et al. 2022; Ahmad et al. 2017), and STAT3 is elevated in rodent models of autism induced by propionic acid (Khera, Mehan, Bhalla, et al. 2022). Our studies provide new mechanistic insights, suggesting that aberrant JAK/STAT signaling in astrocytes may be a key pathway contributing to IL-6-mediated neuronal dysfunction in ASD.

In summary, our findings identify SETD5 as a key regulator of astrocytic homeostasis, whose deficiency is sufficient to drive a reactive astrocyte phenotype characterized by IL-6 accumulation. This response can be attenuated by JAK1/2 inhibition, suggesting that SETD5 loss-of-function dysregulates the JAK/STAT signaling pathway. Future studies using advanced models such as cortical organoids and *in vivo* Setd5+/- mouse models will be essential to further define the physiological relevance of these mechanisms and their contribution to neurodevelopmental pathology.

## Material and Methods

### hiPSC generation and culture

Huma induced pluripotent stem cells (hiPSCs) derived from a neurotypical individual (WT83) or from a subject with non-syndromic autism spectrum disorder (ASD) carrying a missense mutation (I298M) in the SET domain of the *SETD5* gene (*SETD5*^Mut^) (**Suppl Fig 1**) were previously established and characterized (Russo et al. 2018; Marchetto et al. 2017). For the present study, additional independent hiPSCs lines were reprogrammed from *SETD5^I298M^* fibroblasts to establish at least three hiPSC clones. Established Sendai-based reprogramming protocols were used (Priscilla D. Negraes et al. 2021). We generated isogenic CRISPR-corrected *SETD5^Res^*hiPSCs through Applied StemCell (Milpitas, CA) to correct the *SETD5* missense mutation to reference sequence (ATG>ATA). During the process, silent mutations were introduced in heterozygosis 4 nt downstream of corrected mutation (**Suppl. Fig1**). *SETD5* KO heterozygous hiPSCs were generated using two gRNAs; one targeting the first exon, and the other one exon 16 (**Suppl. Fig. 1**). Briefly, hiPSCs were passaged as clumps and maintained for one week before nucleofection. One million cells dissociated with Accutase were resuspended in dPBS. Ribonucleoprotein (RNP) was assembled by mixing pair of gRNA (ATAAUUAUGGGACCACUCAG and CCCAAACACTACAUUCGCUU, 200 pmol each, Synthego), spCas9 protein (40 pmol, Synthego) and incubating at RT for 10 min. Cell suspension, assembled RNP, and 1µg pMax GFP plasmid (Lonza) were then combined and transferred to a cuvette, and electroporated in the NEPA21 system with default setting. Cells were then recovered into pre-warmed mTeSR1 media supplemented with 1x CloneR (Stemcell Technology) and plated on a matrigel-coated 60 mm dish and cultured for four days. Cells were then detached as single cells using Accutase and GFP-expressing cells were sorted with a fluorescence-activated cell sorter (FACS). Approximately 5,000 GFP+ cells were plated in a 10 cm dish coated with Matrigel and cultured with mTeSR1 medium supplemented with CloneR. The following day (24 hours), medium was switched to mTeSR1. One week after plating, single colonies were isolated under an EVOS microscope and expanded. Genomic DNA was isolated using QuickExtract DNA Extraction Solution (LGC Biosearch Technologies) following manufacture’s instructions. PCR genotyping was performed using primers flanking each gRNA (TTGTATCTCAGGTGCCTGCC, TCAGGGAAGTGCAAAAGGGG, CTGCTGCATCTCCTCCTGTC, AACTCACATGGGCAGAGTGG). PCR products were separated on agarose gels to identify clones carrying large deletions. Candidate clones were purified using a PCR purification kit (Qiagen) and subjected to Sanger sequencing for validation of editing outcomes.

For passaging, hiPSCs were picked manually every 5-7 days, seeded onto feeder-free Matrigel-coated dishes (BD Biosciences, San Jose, California), and expanded and maintained with media changes every other day using mTeSR1 or mTeSR plus complete media (StemCell Technologies), and cryopreserved with CryoStor (StemCell Technologies).

### Genomic DNA isolation, genotyping, and karyotyping

Genotypes of all hiPSCs used in this study were confirmed by Sanger sequencing (**Suppl. Fig. 1)**. For the CRISPR-edited *SETD5^Het^*hiPSCs, genomic DNA was extracted with PureLink Genomic DNA Mini kit according to the manufacturer’s instructions. PCR was performed using two sets of primers, one flanking exon 4 and 5 to capture the WT allele and another flanking exon 4 and 16 to capture the deletion (**Table 1**), using the Takara PrimeSTAR Max Enzyme and its suggested PCR settings. PCR reactions were run in agarose gels to confirm a single band amplicon and sent unpurified for Sanger sequencing to Plasmidsaurus. For control, *SETD5^Mut^,* and *SETD5^Res^*cells were lysed and genomic DNA was extracted as previously described (Tenreiro et al. 2025). Briefly, cells were lysed overnight at 56 °C with 100 mM NaCl, 10 mM Tris pH 8.0, 25 mM EDTA pH 8.0, 0.5% v/v SDS, with 0.2 μg/μL Proteinase K (Invitrogen, EO491), and genomic DNA was isolated using conventional phenol:chloroform:isoamyl alcohol (Sigma-Aldrich, P2069-100ML) extraction. For genotyping, 15 ng of genomic DNA was used for PCR amplification using primers in exon 9 flanking the *SETD5* missense mutation with Xpert Fast Hotstart DNA polymerase (Xpert Fast Hotstart Mastermix (2X), grisp, GE35.5001), with the following parameters: 95^◦^C for 3 min, followed by 35 cycles of 30 s at 95^◦^C, 30 s at Tm, and 30 s at 72^◦^C, and then a final step at 72^◦^C for 5 min. PCR amplicons were cleaned up and send for Sanger sequencing (Azenta). Chromatograms were visualized and analyzed with SnapGene Viewer. In SETD5^Het^ clones 7A and 1G, one allele carried a large CRISPR-induced deletion between the exon 4 and exon 16 of the *SETD5* gene, and the other allele contained a single-nucleotide T insertion that shifts the reading frame relative to the first annotated start codon. Sequence inspection indicated that a second downstream start codon remained intact and in-frame with the C-terminal coding region containing the SETD5 domain. On this basis, these lines were classified as heterozygous rather than complete knockout clones.

**Table 1.**
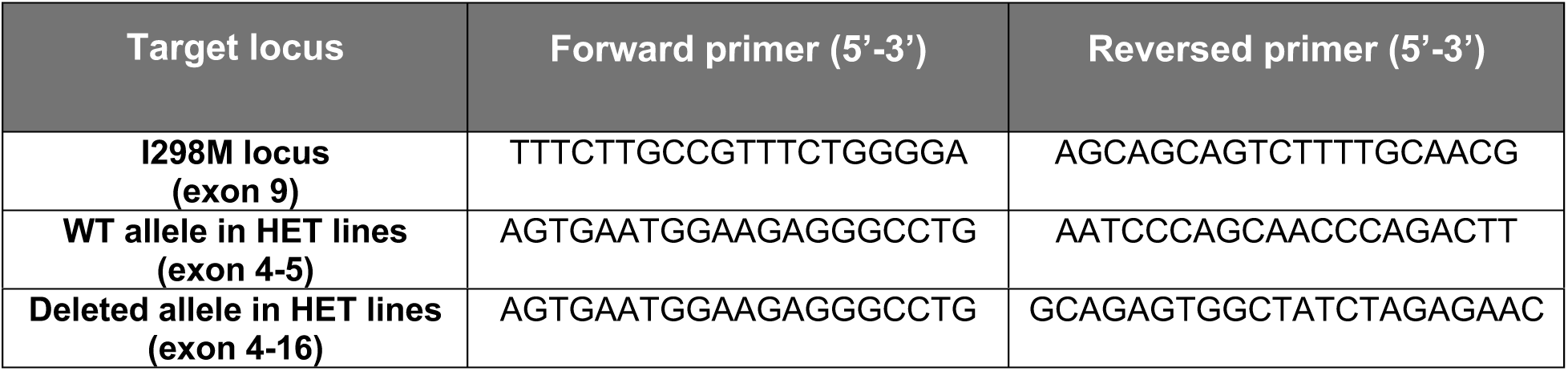
Primer sequences used for genotyping *SETD5*-edited iPSC lines.

karyotype assessment was performed using bead array conducted at the UC San Diego Institute for Genomic Medicine (IGM) (**Suppl Fig. 1**). Pluripotency of all hiPSC lines used in this study was confirmed by immunostaining with well-established pluripotency markers (**Suppl Fig. 1)**.

### Neural induction and astrocyte differentiation

Neuronal progenitor cells (NPC) were generated from hiPSC as previously described (Russo et al. 2018). Briefly, hiPSC were cultured on matrigel-coated dishes in mTeSR1^TM^ medium (StemCell Technologies) until confluency. Then, media was substituted by N2 medium (DMEM/F12 supplemented with N2 (Life Technologies), 1 mM dorsomorphin (R&D Systems), and 1 mM SB431542 (Stemgent) for 24-48 hours. hiPSC colonies were manually lifted and cultured in suspension in N2 media on a rotational shaker (95 r.p.m. at 37°C) for 8 days to form floating embryonic bodies (EBs). EBs were dissociated mechanically by pipetting, plated on matrigel-coated dishes with N2B27 medium (DMEM/F12 supplemented with 0.5x N2, 0.5x B27, 1% penicillin/streptomycin (P/S) (Life Technologies), and 20 ng/mL of FGF-2 (Life Technologies). Emerging neuronal rosettes (easily distinguished under the microscope) were manually picked under EVOS microscope localized inside biosafety hood, dissociated with accutase, and plated onto poly-ornithine (10 ug/mL) /laminin (5 ug/mL) coated dishes. NPCs were fed with N2B27 medium every other day until they reached confluency. To start neuronal differentiation, FGF-2 was withdrew from N2B27 media of confluent cultures of NPCs and media was replaced twice a week for six weeks.

Astrocytes were generated from NPCs as described (Russo et al. 2018) with minor modifications. Briefly, NPCs were incubated with PBS for 5 min in the incubator (37°C). The media was changed, and the cells were lifted with a cell lifter and transferred to low-adherent plates and maintained in suspension in a rotational shaker at 95 r.p.m. at 37°C for 2 days to form floating neurospheres in N2B27 added of FGF (20 ng/mL) media. After 2 days, and to start differentiation, the media was changed for N2B27 supplemented with Rock inhibitor (RI) (5 μM) was added and kept for 2 days in this media. Then, N2B27 media was substituted by Astrocytic Growth Medium AGM^TM^ (Lonza) and the neurospheres/astrospheres were kept in for 2 weeks in suspension. Then, the astrospheres were plated onto poly-ornithine/laminin-coated plates to allow cells to migrate out of the astrospheres and passaged after one month with Papain and DNase (Worthington Biochemical Corp). For experiments, astrocytes were used after passages 3-4 without including spheres. Spheres were kept in a parallel dish for the continuous generation of astrocytes.

### Immunofluorescence staining

Cells were fixed with 4% paraformaldehyde in PBS for 15 min at room temperature, washed three times with PBS, permeabilized with 0.1% Triton X-100 for 15 min, and blocked with 2% BSA in PBS with 0.1% Triton X-100 for 1 hour at room temperature followed by incubation with primary antibodies diluted in 2% BSA in PBS with 0.1% Triton X-100 (chicken anti-GFAP, Abcam Ab4674, 1:1000; rabbit anti-AQP4, Abcam Ab46182, 1:500; mouse anti-S100B, Abcam Ab14849, 1:500; chicken anti-MAP2, Abcam Ab5392, 1:1000; mouse anti-VGLUT1, Synaptic Systems 135311, 1:500; rabbit anti-HOMER1, Synaptic Systems 160003, 1:500; rabbit anti-SOX2, Cell Signaling 2748, 1:500; mouse anti-Nestin, Abcam Ab22035, 1:500; goat anti-Nanog, Abcam Ab77095, 1:500; rabbit anti-LIN28A, Cell Signaling 3978S, 1:500) overnight at 4°C. Next day, cells were washed three times with PBS and incubated with secondary antibodies diluted in 2% BSA in PBS with 0.1% Triton X-100 (Alexa Fluor 488, 555 and 647, Life Technologies, 1:2,000) for 1 hour at room temperature. After, cells were washed three times with PBS and followed by incubation with DAPI (1:10,000) for 5 min. Slides were mounted using ProLong Gold Antifade Mountant (Life Technologies). Images were obtained using Zeiss fluorescence microscope equipped with Apotome (Axio Observer Apotome, Zeiss).

### Astrocyte morphology quantification

After fixing the astrocytes and performing an immunofluorescence protocol, we imaged the cells using the Axio Observer, Zeiss. The GFAP staining was used to classify the cell morphology, which was analyzed with ImageJ2 2.14.0/1.54f, primarily as maximum intensity projections. Morphology was classified into three categories: fibroblast-like, arborized, and polarized cells, with analysis of >100 cells from three batches at × 20 magnification as previously reported (Jones et al. 2017).

### Reactive oxygen species (ROS) measurement

Intracellular levels of ROS were quantified using luminescence assays ROS-Glo (Promega), and performed according to the manufacturer’s instructions. Briefly, 4 x 10^4^ hiPSC-derived astrocytes were plated in each well of a white 96-well plate (Corning) and maintained until cells reached ∼80% confluence. Then, the media was changed, and measurements were performed 24 hours later. Six wells per genotype from the same astrocytic differentiation were quantified. An extra set of four wells was used to assess cell viability by measuring ATP levels using the CellTiter-Glo Luminescent Cell Viability Assay (Promega), according to the manufacturer’s instructions.

### Extracellular glutamate quantification

4 x 10^4^ hiPSC-derived astrocytes were plated per well in a 96-well plate and cultivated until they reached ∼80% confluence. Then, the media was changed by fresh media and 24 hours later colorimetric Glutamate assay kit (Promega) was performed following the manufacturer’s recommendations. This assay was done in triplicate and normalized by numbers of viable cells obtained using CellTiter-Glo Luminescent Cell Viability Assay (Promega).

### Extracellular cytokines/chemokines quantification

hiPSC-derived astrocytes were dissociated, and 1 x 10^4^ cells were plated per well in 384-well coated-plated and maintained until they reached 70-80% confluency. Then, the media was removed, fresh media was added and 24 h later, 25 μl of media was collected to measure levels of IL1-β, IL-8, IL-10, IL-12p70, IL-6, and TNF using the human inflammatory cytokine kit (BD Biosciences) following the manufacturer’s instructions using a fluorescence-activated cell sorting (FACS) Canto 2 instrument (BD Biosciences).

### Synaptic puncta quantification

We used immunofluorescence staining to count co-localizing presynaptic VGlut1 and postsynaptic Homer1 immunoreactive spots along dendrites marked by MAP2 as previously described (Russo et al. 2018). Synaptic puncta was visualized and quantified over 50 µm segments of MAP2-positive dendrites, using a Z1 Axio Observer Apotome fluorescence microscope (Zeiss).

### Neurite length measurement

We conducted the morphological analysis of neurons utilizing the Neurolucida Neuron Tracing Software (MBF Bioscience, version 2017.01.1). iPSC-derived neurons were transduced with Synapsin1 promoter driven EGFP expression, aged 8 weeks, and stained for CTIP2 (indicating their identity as layer V/VI neurons). Only those neurons that had nuclei stained with CITP2 and had neurites with uniform EGFP staining throughout were analyzed. Neurites were classified as dendrites if they met specific criteria such as progressively diminishing thickness from the soma, branching at acute angles, and the presence of dendritic spines. Neurolucida v.9 software (MBF Bioscience, Williston, VT) on a Nikon Eclipse E600 microscope with a 40× oil immersion lens was used to collect images. in three dimensions. Analysis did not differentiate between apical and basal dendrites, aggregating the lengths of all neurites/dendrites per neuron.

### Neuron co-cultures with astrocytes or astrocyte conditioned media

hiPSC-derived neurons/astrocytes co-culture experiments were performed as described (Russo et al. 2018). Briefly, six-week differentiated neuronal cultures were dissociated using Accutase and were resuspended in MACS buffer and incubated with phycoerythrin (PE)-conjugated CD184 and CD44 antibodies (BD Biosciences) at 4°C for 15 minutes in the dark. Cells were washed and spun down at 300xg, and the cell pellet was resuspended at the ratio of 80 ml/10^7^ cells and added 20 ml of magnetic beads coated with anti-PE antibodies (BD Biosciences) and incubated for 15 min at 4°C. 2 x 10^4^ enriched neurons after magnetic cell sorting were plated on top of hiPSC-derived astrocytes cultured at ∼80% confluency and kept with AGM for two weeks. Then, cells were fixed and processed for immunofluorescence as described above. Synaptic puncta and neurite length were quantified and measured, respectively, as explained above.

For conditioned media experiments, astrocytes were plated on matrigel-coated 10 cm dishes in AGM Astrocyte Growth Medium (Lonza). Vehicle or CYT387 was added to the astrocytes and 24 hour later media was collected, sterile-filtered, aliquoted and stored in −80 before use.

NPCs were differentiated into neurons in matrigel-coated 10 cm dishes in Neurobasal + B27 media for differentiation for 21 days. Then, neurons were replated at low density (50,000/cm2) in PDL and laminin-coated glass chamber slides (8 chambers, LabTekII) in Neurobasal + B27. 24 hours later, media was changed to the astrocyte conditioned media described above. Half media changes with conditioned media were performed every other day for 2 weeks before cells were fixed with PFA for immunofluorescence imaging. Neuron images were traced and quantified with ImageJ.

### Transcriptomic analyses and gene expression profile for *SETD5* gene

To assess the expression profile of *SETD5*, we analyzed the available bulk RNA-sequencing datasets generated from human fibroblasts, hiPSCs, and hiPSC-derived NPC, neurons, and astrocytes (P. D. Negraes et al. 2017; Chailangkarn et al. 2016; Mesci et al. 2024) and from unpublised datasets) derived from neurotypical individuals. Raw sequencing reads were first subjected to quality control using NGS QCToolkit (Patel and Jain 2012) to assess read quality, adapter contamination, sequence duplication, and overall sequencing performance. After quality assessment and preprocessing, reads were aligned to the human reference GRCh38 genome using the STAR aligner (Dobin et al. 2013). STAR was used because it is optimized for spliced RNA-seq alignment and allows accurate mapping of reads across exon–exon junctions. Aligned reads were used to generate gene-level count matrices based on annotated genomic features using the HTSeq framework (Anders et al. 2015). Gene-level read counts were then imported into DESeq2 (Love et al. 2014) for normalization and differential expression analysis. Raw count data, rather than normalized expression values, were used as input for the statistical model. DESeq2 normalization was performed using size factors estimated by the median-of-ratios method, which corrects for differences in sequencing depth and RNA composition across samples. For visualization purposes, expression values were extracted as DESeq2 size factor–normalized counts.

Differential expression analysis was performed using DESeq2 (Love et al. 2014). Cell type was considered as the main experimental variable, and pairwise contrasts were performed using astrocytes as the reference group that was compared to other cell types: fibroblasts, hiPSCs, NPCs and neurons. For each comparison, DESeq2 estimated the log2 fold change, standard error and FDR (adjusted p-value). Statistical significance was assessed using the Wald test implemented in DESeq2. Correction for multiple testing was performed using the Benjamini–Hochberg false discovery rate method across the genes included in the transcriptome-wide analysis. Genes were considered statistically significant when the FDR was below 0.05. For the gene *SETD5*, DESeq2 results were extracted from each pairwise comparison. Mean normalized expression values were also calculated for astrocytes and each compared cell type.

For visualization, gene expression plots were generated in Python using the size factor-normalized counts obtained from DESeq2. The normalized expression values for *SETD5* gene were organized by cell type, with each point representing one biological replicate. Cell type labels containing replicate suffixes (sequential numbers) were collapsed into their corresponding biological group names. Violin plots were generated using the seaborn Python package to show the distribution of normalized expression values across cell types. The y-axis represents the expression levels (as size factor–normalized counts) to clarify that these values correspond to DESeq2-normalized rather than raw counts. The statistical annotations shown on the violin plot correspond to the DESeq2 FDR-adjusted p-values obtained from the transcriptome-wide differential expression analysis.

### Single cell RNA-seq

Monolayers of hiPSC-derived astrocytes were washed with DPBS and dissociated with Accutase mixed with 10:1 papain (Worthington, LK003176) and DNaseI (Worthington, LK003170) dissolved in Earle’s Balanced Salt Solution (EBSS; Worthington, LK003188) for 10 or 30 min at 37 °C and 5% CO_2_. Accutase/Papain mix was inactivated with Astrocytic Growth Medium and filtered with a 70-μm strainer. Cell count and viability was determined using ChemoMetec cell counter or a hematocytometer. The minimum population viability threshold for downstream scRNA-seq processing was set at 80%. Cells were pelleted for 3 min at 100 x g at 4°C and resuspended in cold 0.1% BSA in DPBS.

scRNA-seq libraries were constructed using the Chromium™ Single Cell 3’ v3 Library kit (10x Genomics (Zheng et al. 2017)) according to manufacturer descriptions. Approximately 15,000 cells were loaded per sample. Reverse transcription and other amplification steps were carried out on a T100 thermal cycler (Bio-Rad). After reverse transcription, GEMs (Gel beads in the emulsion) were lysed, and cDNA was cleaned up with MyOne Silane Beads (Thermo Fisher Scientific). Next, single-stranded cDNA was PCR-amplified for 12 cycles and purified using SPRIselect Reagent Kit (Beckman Coulter). Next, cDNA was enzymatically fragmented, followed by double size selection with SPRIselect Reagent Kit (B23317, Beckman Coulter). Subsequently, adaptors were ligated, and libraries were constructed using PCR. Another round of double size selection was performed using SPRIselect Reagent Kit to generate final libraries with a size of 200-700bp. Final libraries were quantified using Qubit® dsDNA HS Assay Kit (Thermo Fisher Scientific) and size distribution was measured using Tapestation (High Sensitivity D1000, Agilent). Average fragment size of successful libraries was 500 bp. The libraries were loaded at a concentration of 13 pM and sequenced on a Hiseq 4000 sequencer (Illumina) with the following parameters (Read1 26 cycles; Index 1 8 cycles; Read 2 98 cycles). Eight sequencing libraries were analyzed, comprising four genotypes: JB_70, JB_635 (controls or Reference); JB_71, JB_638 (SETD5^Mut^); JB_636, JB_637 (SETD5^Het^); and JB_639, JB_72 (SETD5^Res^). Data preprocessing and analysis were conducted using several computational tools, primarily Seurat, CellBender, DoubletFinder, and Harmony, with additional pathway enrichment analysis using EnrichR. We used the CellBender tool to eliminate ambient and background RNA from the count matrix generated by CellRanger, setting the false positive rate (FPR) at 0.05, using the “remove-background” function. Quality Control (QC) and Filtering were applied for each dataset that underwent QC, with filtering parameters set based on nFeature_RNA and the percentage of mitochondrial reads (percent.mt), which is a tool that maps the mitochondrial genome. The filtering parameters varied across samples: - JB_70: 20 < nFeature_RNA < 8000; percent.mt < 25. JB_635: 200 < nFeature_RNA < 6500; percent.mt < 10. JB_71: 20 < nFeature_RNA < 8000; percent.mt < 30. JB_638: 200 < nFeature_RNA < 4500; percent.mt < 15. JB_636: 200 < nFeature_RNA < 6000; percent.mt < 10. JB_637: 200 < nFeature_RNA < 6000; percent.mt < 15. JB_639: 200 < nFeature_RNA < 7000; percent.mt < 20. JB_72: 200 < nFeature_RNA < 7500; percent.mt < 25. The DoubletFinder was applied to the CellBender output files (.h5) for doublet identification and removal. Initial creation, clustering, and normalization of the first eight Seurat Objects were achieved via SCTransform. DoubletFinder parameters included PCs 1:15, pN at 0.25, and dataset-specific pK values determined using the “find.pK()” function. The parameter nExp was estimated based on the “Estimated Number of Cells” from the 10X Cell Ranger output. The “DF.classifications” column in the metadata was used to identify and remove doublets. Data Integration and Normalization: Post-doublet removal, the datasets were merged using Seurat’s merge() function into a single Seurat object, which was then normalized using SCTransform. Batch Correction and Clustering Analysis: Harmony was utilized for batch correction. Clustering analysis was conducted with a resolution of 0.2, employing the first 10 principal components. Differential Gene Expression Analysis and Pathway Enrichment: Differential gene expression across clusters was determined using the FindAllMarkers function in Seurat. Gene Ontology (GO) pathway enrichment analysis was performed using the GO_Biological_Process_2023 gene set in EnrichR.

### Single nuclei ATAC-seq

hiPSC-derived astrocytes were dissociated as above. Threshold to proceed was at least 80% viable cells and >500,000 cells. Cells were then centrifuged (300 xg, 5 min, 4°C, ramp speed 3/3) in a swinging bucket centrifuge. Supernatant was removed and cells were resuspended in 100 μl of lysis buffer consisting of 10 mM Tris-HCl pH 7.4 (15567-027, Invitrogen), 10 mM NaCl (AM9760G, Invitrogen), 3 mM MgCl_2_ (AM9530G, Invitrogen), 0.1% Tween-20 (P7949-100ML, Sigma Aldrich), 0.1% IGEPAL CA-630 (I8896-50ML, Sigma Aldrich), 0.01% Digitonin (G9441, Promega), 1% BSA (7500804, Lampire Biological) in PBS (21-040-CV, Corning) in nuclease free water (46-000-CV, Corning). Cells were incubated for 4 minutes on ice. 1 mL of wash buffer consisting of 10 mM Tris-HCl pH 7.4, 10 mM NaCl, 3 mM MgCl_2_, 1% BSA in PBS, and 0.1% Tween-20 in nuclease free water was then added to nuclei and mixed 5 times. Nuclei were then centrifuged (500 xg, 5 min, 4°C, ramp speed 3/3) in a swinging bucket centrifuge. Supernatant was removed and cells were resuspended in nuclei buffer consisting of 1X Nuclei Buffer (PN-2000153, 10x Genomics) in nuclease free water. Nuclei were stained with Trypan Blue and counted with a hemacytometer.

scATAC-seq libraries were generated using kits from 10x Genomics per manufacturer specifications (PN-1000175, 10x Genomics). Tagmentation was performed on 16,000 nuclei and which were then loaded for GEM capture. Libraries were prepared for index PCR and indexed using the single index plate (PN-1000212, 10x Genomics) and 10 PCR cycles. Final libraries were cleaned up using SPRISelect reagent (PN-B23319, Beckman Coulter). Library QC was performed with Qubit High Sensitivity Assay (Q32851, Invitrogen) and Tapestation D1000 (5067-5882 and 5067-5883, Agilent). Libraries were sequenced on a NextSeq500 (Illumina) for 10-20 million reads to assess data quality, and then on a a NovaSeq 6000 (Illumina) for a target of 300 million reads. Sequencing reads were demultiplexed to fastqs using cellranger mkfastq (10x Genomics) and filtered, mapped, and aligned to cell barcodes using cellranger-atac v 2.0.0 (10x Genomics).

Following demultiplexing, scATAC-seq was analyzed using SnapATAC 2 (K. Zhang et al. 2024). Cells with less than 1,000 ATAC fragments or a TSS enrichment score of less than 7 were filtered out, while the top 50,000 most accessible features were selected for downstream processing. Likely doublets were removed using the scrublet method. Peaks were called using MACS3 (Y. Zhang et al. 2008), merged, and differential peak testing performed using default parameters. Differential peaks were associated with genes using HOMER (Heinz et al. 2010) and scanned for motifs using SnapATAC2. Motif enrichment was calculated with the motif_enrichment function from SnapATAC2 using the hypergeometric method. Functional enrichment of the associated protein-coding genes was performed using Metascape (Y. Zhou et al. 2019).

### Epigenetic compound library screening in SETD5^Mut^ astrocytes

*SETD5^Mut^* hiPSC-derived astrocytes were dissociated and plated at 2 x 10^4^ cells per well in coated 384-well plates. Four wells were seeded with control astrocytes. Cells were maintained until they reached approximately 80% confluency. The medium was then replaced, and 24 hours later, extracellular IL-6 and IL-8 levels in the supernatants from 4 control wells and 12 SETD5^Mut^ wells were measured using BD™ Cytometric Bead Array (CBA) Human IL-6 or IL-8 Flex Set and a FACSCantoII -V96300991. Plates were used for compound screening only if extracellular IL-6 and IL-8 levels in the SETD5^Mut^ wells were at least twofold higher than those in the control wells.

384-well plates containing a library of approximately 350 compounds targeting epigenetic enzymes were assembled in-house by the National Center for Advancing Translational Sciences (NCATS). The library included inhibitors of histone deacetylases (HDACs), sirtuins (SIRTs), lysine demethylases, histone acetyltransferases (HATs), DNA methyltransferase (DNMTs), as well as SIRTs activators and control compounds. Compounds were tested at final concentrations of 1, 10, and 20 μM. Compound plates were thawed at room temperature for 40 min, centrifuged at 1,000 rpm for 1 min, and resuspended in 60 μl of AGM per well. Subsequently, 50 μl from each well was transferred to the corresponding well containing seeded SETD5^Mut^ astrocytes while maintaining the original plate layout. Astrocytes were treated for 24 h, and 25 μl of media was collected to measure levels of IL-6 and IL-8. Once the culture medium was removed, cell viability was assessed by measuring ATP levels using the CellTiter-Glo Luminescent Cell Viability Assay (Promega). IL-6 and IL-8 levels were measured using BD™ Cytometric Bead Array (CBA) Human IL-6 or IL-8 Flex Set in a FACSCantoII -V96300991. IL-6 and IL-8 standard curves were generated following manufacturer’s instructions. Wells with levels of IL-6 or IL-8 below or above the standard range were discarded. IL-6 and IL-8 levels were normalized to ATP levels, as a surrogate for viable cell number. Secondary and validation experiments were performed similarly using independently generated cultures of hiPSC-derived astrocytes.

## Acknowledgments

We acknowledge K. Jepsen at the UCSD Institute of Genomic Medicine funded by the National Institutes of Health (NIH) grant (#S10 OD026929) and NIEHS (R01 #ES027981) to BL.

## Author Contributions

Conceptualization: ARM, IRF, JSS, PM, AAQ

Experimental investigation: AAQ, IRF, JSS, AS, XS, GC, DP, KD, LBVC, LLSN, MFT, LFPB

Data analysis: AAQ, IRF, JSS, AS, GC, KD, MC, AF, BT, MFT, LFPB, ED, PM, RHH, AW, IGB.

Manuscript writing – AAQ (Original Drafts: JSS & IRF). Reviewed and editing: ARM, AG, CB, IGB, JM, MF

Funding acquisition: ARM, AAQ, AG, CB

Project administration and oversight: ARM and AG

---

**Supplementary Figure 1.**
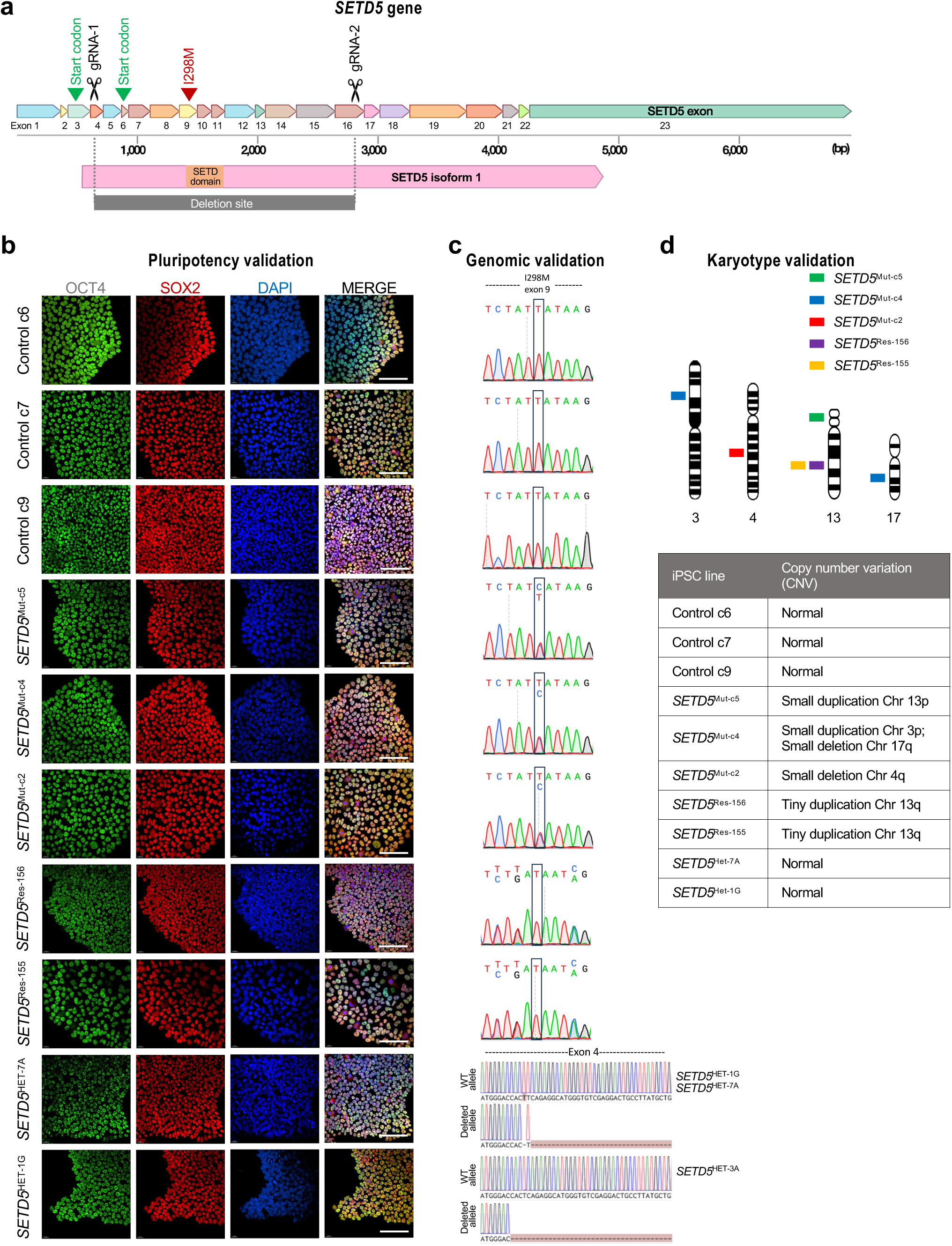
Cellular and genomic characterization of iPSC lines. **a.** Schematic of the CRISPR-Cas9 strategy used to generate heterozygous deletions in SETD5. sgRNAs targeting exon 4 and exon 16 were used to induce a large deletion spanning the intervening region. Two annotated translation start sites are indicated. **b.** Representative immunofluorescence images of control and genome-edited iPSC lines, including SETD5 mutant, heterozygous, and rescue clones, stained for pluripotency markers OCT4 (green) and SOX2 (red), with nuclei counterstained with DAPI (blue). Scale bar, 100 μm. **c.** Sanger sequencing and allele alignments confirming SETD5 genotypes in edited hiPSC lines. In heterozygous clones HET-7A and HET-1G, one allele carries a large deletion between the sgRNA target sites, whereas the second allele contains a single-nucleotide T insertion. The insertion is predicted to shift the reading frame relative to the first annotated start codon; however, a downstream start codon remains present. **d.** Copy number variation (CNV) analysis of iPSC lines. Ideograms indicate chromosomal regions with detected CNVs in individual SETD5 mutant and rescue clones. Clone identities (c2, c4, c5, 155, and 156) are indicated by color. CNVs are shown at the chromosomal arm level based on the resolution of the assay. Only chromosomes with detected alterations are displayed. The type of CNV (duplication or deletion) is summarized in the accompanying table.

**Supplementary Fig. 2.**
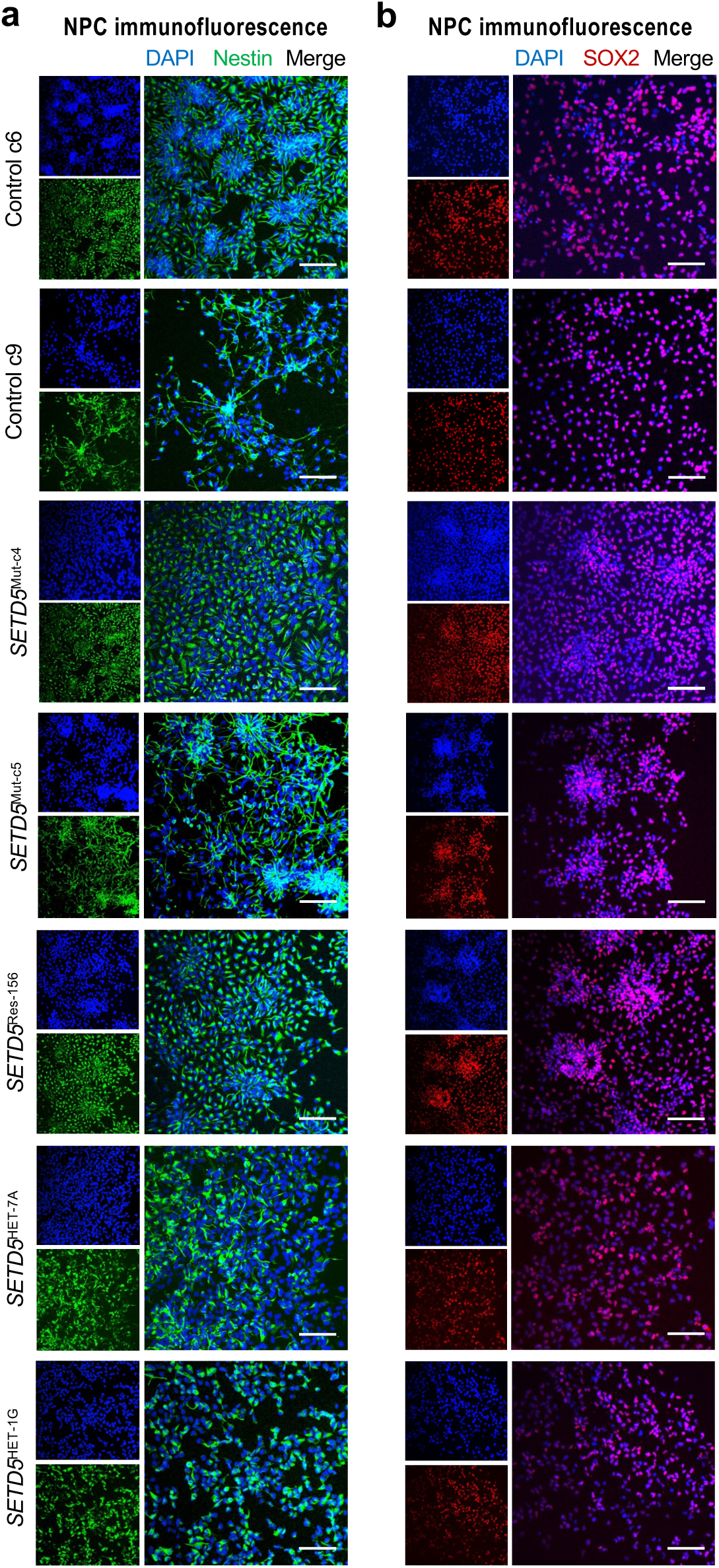
Characterization of neural progenitor cells (NPCs) derived from control and SETD5-edited hiPSC lines. **a.** Representative immunofluorescence images of NPCs derived from control (c6, c9), SETD5 mutant (c4, c5), rescue (156), and heterozygous (7A, 1G) lines stained for the neural progenitor marker Nestin (green) and DAPI (blue). **b.** Representative immunofluorescence images of the same NPC lines stained for the neural progenitor marker SOX2 (red) with DAPI (blue). All lines exhibit expression of canonical NPC markers Nestin and SOX2, confirming successful neural progenitor identity across genotypes. Scale bars, 100 μm.

**Supplementary Figure 3.**
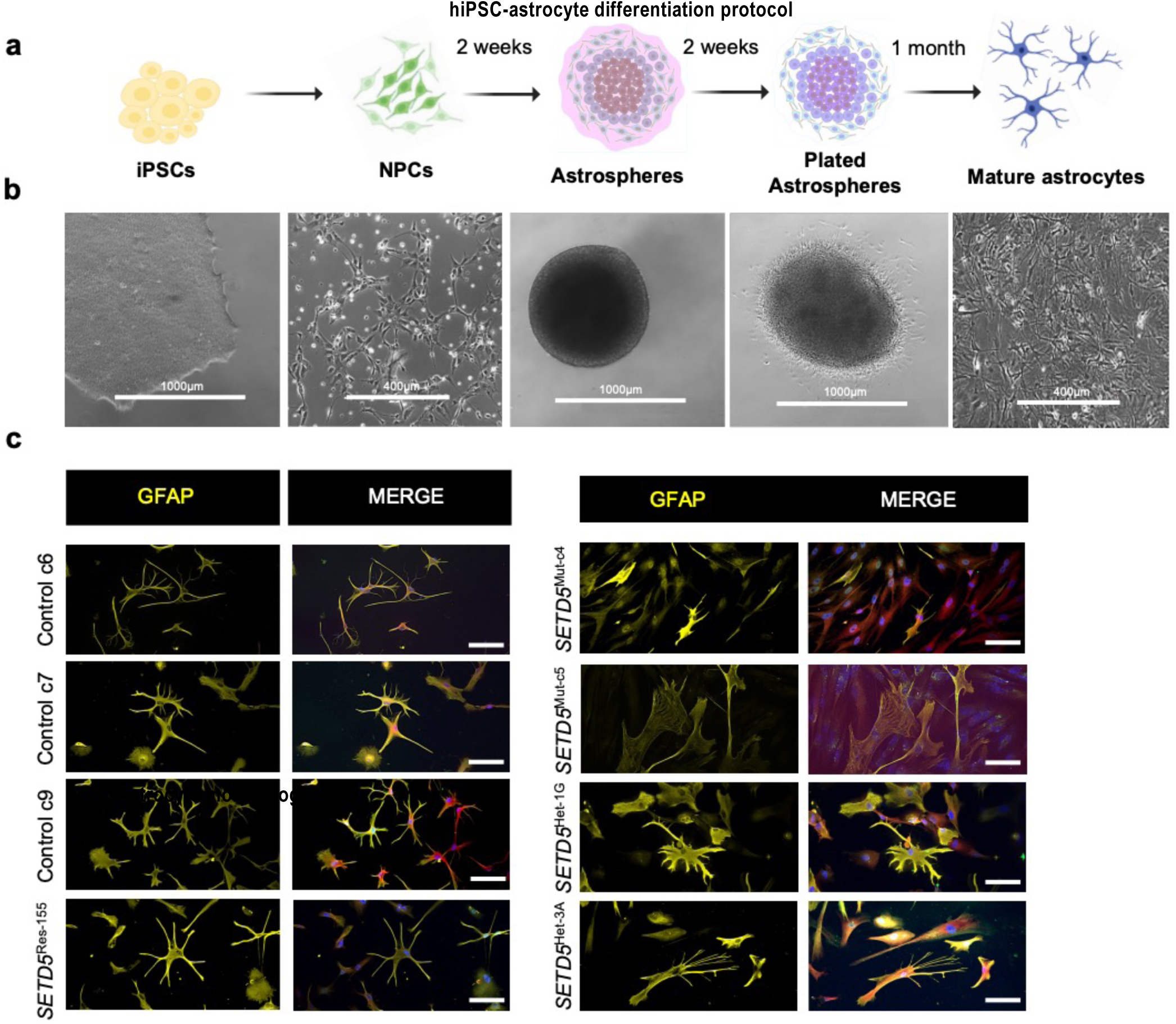
Generation and characterization of human astrocytes from iPSCs. **a.** Schematic overview of the differentiation timeline from human induced pluripotent stem cells (hiPSCs) to neural progenitor cells (NPCs), astrospheres, plated astrospheres, and mature astrocytes. Key stages and approximate durations are indicated. **b.** Representative brightfield images showing morphological progression across differentiation stages. **c.** Immunofluorescence analysis of mature astrocytes from control and SETD5-mutant lines, stained for astrocytic markers GFAP, AQP4, and S100B, with DAPI nuclear staining. Representative merged images illustrate astrocyte morphology and marker distribution across genotypes.

**Supplementary Figure 4.**
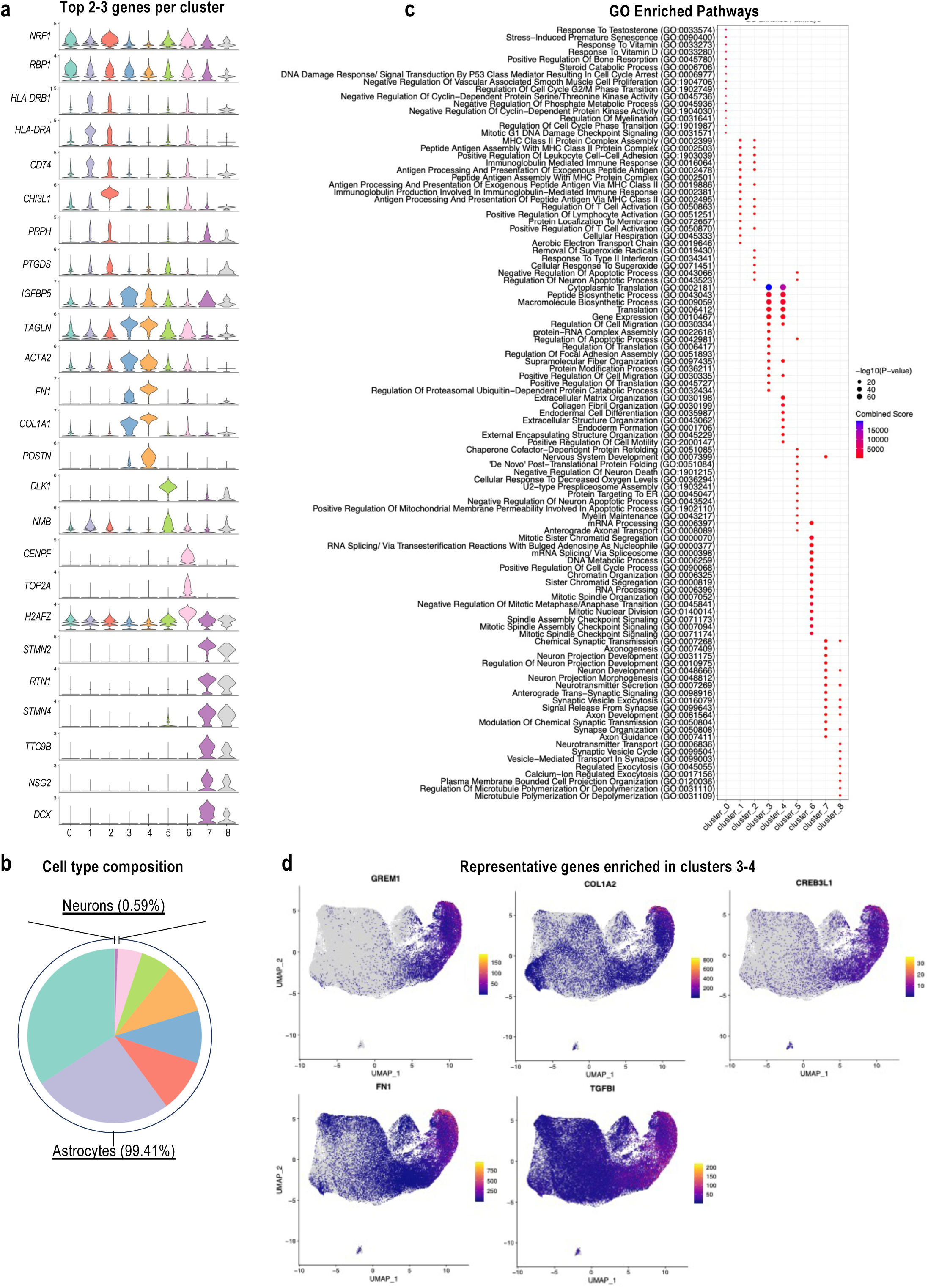
QC of the scRNA-seq datasets. **a.** Violin plots showing expression levels of top 2-3 genes per cluster. **b.** Pie chart representing cell type composition across clusters. **c.** Heatmap of gene ontology (GO) presenting all cluster analyses illustrating the functional enrichment of differentially expressed genes. **d**. UMAP plots showing the spatial distribution of indicated genes across the dataset, with each point representing a single cell.

**Supplementary Figure 5.**
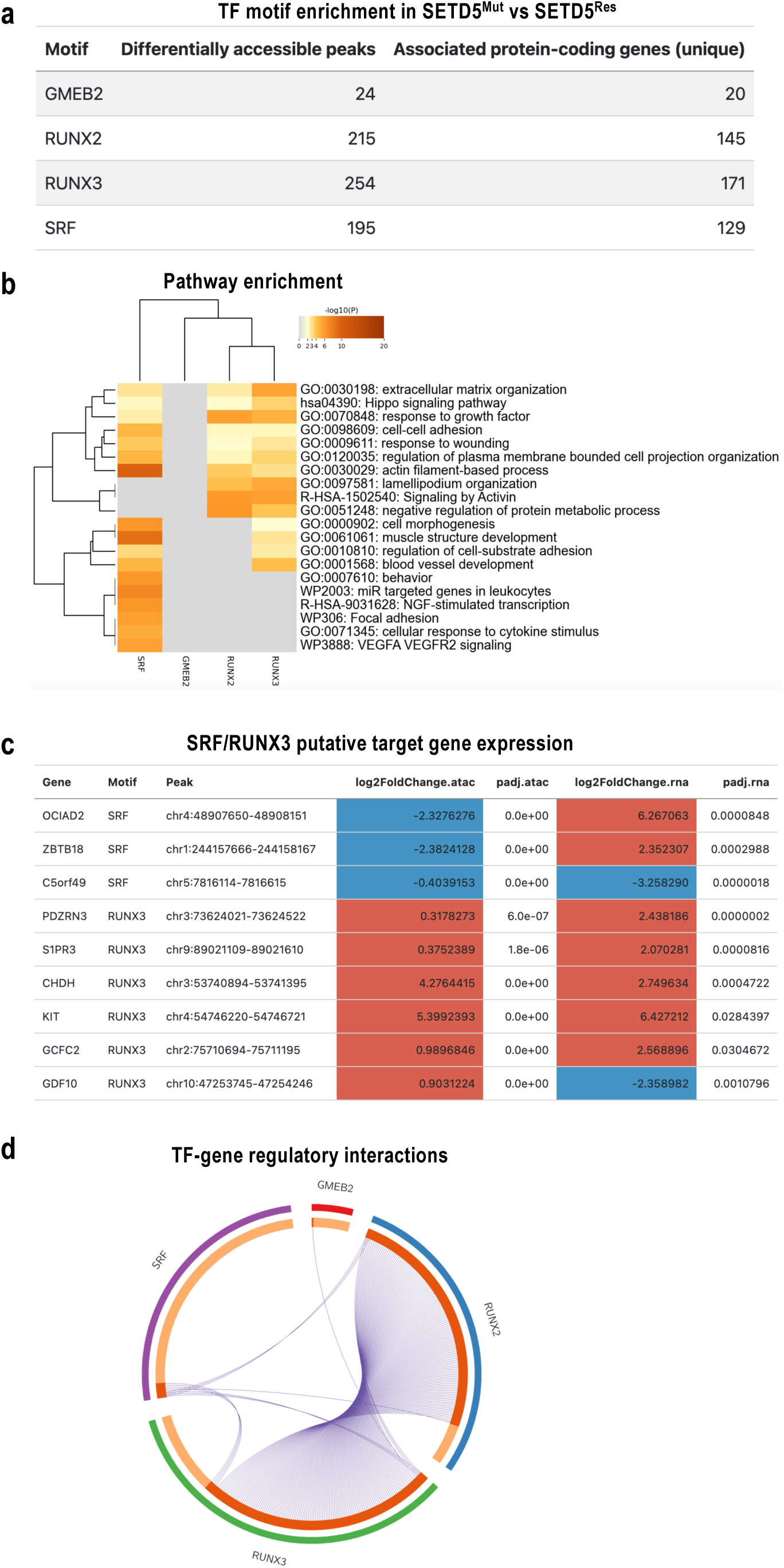
snATAC-seq analysis reveals SETD5^Mut^ transcription factor–specific chromatin accessibility programs. **a.** Motif enrichment analysis of differentially accessible peaks identifies transcription factors associated with chromatin remodeling. The number of peaks containing each motif and the number of unique associated protein-coding genes are shown for GMEB2, RUNX2, RUNX3, and SRF, highlighting strong enrichment for RUNX and SRF family members. **b.** Hierarchical clustering heatmap of gene ontology (GO), KEGG, and Reactome pathway enrichment associated with motif-linked accessible regions. Enriched pathways include extracellular matrix organization, Hippo signaling, cell adhesion, actin cytoskeleton regulation, cytokine response, and VEGFA–VEGFR2 signaling, indicating roles in cell structure, signaling, and inflammatory responses. **c.** Integration of snATAC-seq and gene expression data showing representative genes associated with SRF and RUNX3 motifs. The table displays genomic peak locations, chromatin accessibility changes (log2 fold change, ATAC), and corresponding transcriptional changes (log2 fold change, RNA), with adjusted p-values, illustrating concordant and discordant regulatory relationships. **d.** Circos plot depicting predicted regulatory interactions between transcription factor motifs (GMEB2, SRF, RUNX2, RUNX3) and their associated target genes, based on co-occurrence of motifs within differentially accessible peaks. The density of connections highlights extensive regulatory networks, with RUNX and SRF motifs showing the highest connectivity. Inner colored arcs represent red=shared target genes, orange=TF-specific target genes.

**Supplementary Figure 6.**
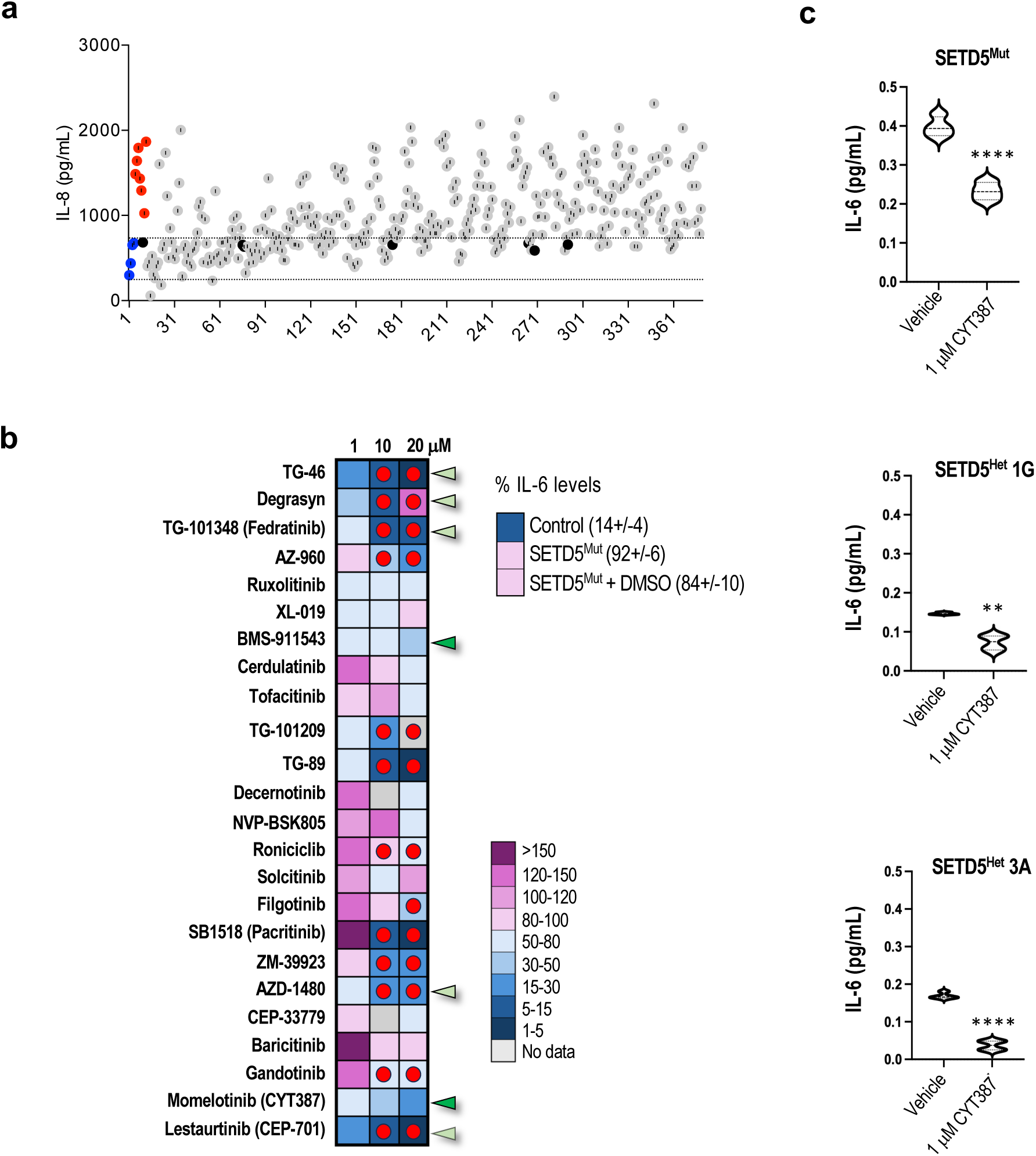
Modulating IL-6 and IL-8 levels in hiPSC-*SETD5^Mu^*^t^ astrocytes. **a.** High-throughput screening results showing the IL-8 levels after treatment with a 350-compound library. **b.** Heatmap representing extracellular IL-6 levels from *SETD5^Mut^* astrocytes treated with the indicated compounds (left) at three different concentrations indicated on top. Extracellular IL-6 levels from Control and *SETD5^Mut^*astrocytes untreated or DMSO-treated were included as controls (top right). Levels are expressed as a percentage of IL-6 levels from untreated *SETD5^Mut^*astrocytes. The color scale indicates ranges of values. Red dots indicate conditions in which cell viability was reduced. **c.** Bar graph presenting extracellular IL-6 levels in *SETD5^Mut^*, and *SETD5^Het^*treated with vehicle or with the JAK1/2 inhibitor CYT387.

