## Supplemental Table 1 for "SETD5 dysfunction in human astrocytes drives IL-6-mediated neuronal impairments via the JAK/STAT signaling pathway"

**Suppl. Table 1** – Expression profile (size-factor normalized counts) of biological replicates of fibroblasts, iPSCs, NPCs, neurons, and astrocytes.

| Cell type (replicate) | Expression |
|---|---:|
| Fibroblast | 5201.05951 |
| Fibroblast.1 | 5536.13352 |
| Fibroblast.2 | 2323.66136 |
| Fibroblast.3 | 4687.80198 |
| Fibroblast.4 | 5733.08036 |
| Fibroblast.5 | 2411.75299 |
| iPSC | 3388.22664 |
| iPSC.1 | 3600.98734 |
| iPSC.2 | 12977.5736 |
| iPSC.3 | 10273.8965 |
| iPSC.4 | 9042.83262 |
| iPSC.5 | 9418.03013 |
| iPSC.6 | 10457.6403 |
| iPSC.7 | 7712.05089 |
| NPC | 12217.1826 |
| NPC.1 | 10748.946 |
| NPC.2 | 10777.2582 |
| NPC.3 | 9826.87187 |
| Neuron | 6351.73727 |
| Neuron.1 | 6779.68711 |
| Neuron.2 | 6933.06269 |
| Neuron.3 | 6836.42388 |
| Astrocytes | 18260.9807 |
| Astrocytes.1 | 17496.2023 |
| Astrocytes.2 | 18062.4059 |
| Astrocytes.3 | 14518.7415 |
| Astrocytes.4 | 14611.6618 |
| Astrocytes.5 | 14370.9041 |
