## Supplemental Table 2 for "SETD5 dysfunction in human astrocytes drives IL-6-mediated neuronal impairments via the JAK/STAT signaling pathway"

**Suppl. Table 2** – Statistical significance of *SETD5* expression in astrocytes compared to fibroblasts, iPSCs, NPCs, neurons. Mean expression values represent DESeq2 size factor–normalized counts for the reference and compared cell types. The log2FoldChange indicates the DESeq2-estimated log2-transformed expression difference between groups (positive log2 fold change values indicate higher expression in astrocytes relative to the compared cell type). The lfcSE represents the standard error of the log2FoldChange estimate. The FDR value corresponds to the Benjamini–Hochberg false discovery rate–adjusted p-value calculated by DESeq2 across the transcriptome-wide analysis.

| Reference cell type | Compared cell type | Mean expression in reference cell type | Mean expression in compared cell type | log2FoldChange | lfcSE | FDR |
|---|---|---|---|---|---|---|
| Astrocytes | Fibroblast | 16220.14937 | 4315.58162 | 1.909817285 | 0.27720573 | 4.44E-11 |
| Astrocytes | iPSC | 16220.14937 | 8358.904735 | 0.956220732 | 0.259169 | 0.000649863 |
| Astrocytes | Neuron | 16220.14937 | 6725.227736 | 1.270134058 | 0.309879757 | 0.000164561 |
| Astrocytes | NPC | 16220.14937 | 10892.56469 | 0.574451128 | 0.3097261 | 0.144457254 |
